# Avocado-derived compounds alter lipid homeostasis and lipid droplets profile in *Caenorhabditis elegans*

**DOI:** 10.64898/2026.08.22.746425

**Authors:** Yamanappa Hunashal, Suma Gopinadhan, Raihanah Harion, Fathima S. Refai, Yasmine Moussa, Liaqat Ali, Kristin C. Gunsalus, Hala Zahreddine Fahs, Gennaro Esposito, Fabio Piano

## Abstract

**Background:** Natural compounds from avocado fruit (avocadene, avocadyne, and acetate derivatives) exhibit notable biological activity, although their molecular mechanisms remain unclear. The avocado-derived lipids exert potent nematocidal activity against several parasitic nematodes. In *Caenorhabditis elegans (C. elegans)*, those compounds caused concentration-dependent toxicity, impairing first stage larval growth, egg hatching, and adult survival. Treated worms exhibited impaired mitochondrial respiration, reduced oxygen consumption, and elevated reactive oxygen species. These effects suggest that avocado lipids disrupt mitochondrial function and lipid metabolism, in part by inhibiting acetyl-CoA carboxylase, the rate-limiting enzyme of fatty acid biosynthesis.

**Methods:** We investigated the effects of these compounds on the lipid profile of *C. elegans* and their association with endogenous lipid pools using NMR spectroscopy, click-chemistry-based fluorescence labeling, thin-layer chromatography (TLC), and microscopy.

**Results:** Lipidomic analysis of stage 4 larvae (L4) and embryos treated with avocadene acetate revealed increased lipid NMR signals. Fluorescence-assisted TLC and NMR further suggested that avocadyne preferentially associates with triglyceride-linked fatty acids, particularly monounsaturated and flexible polyunsaturated chains, without detectable interactions with conformationally-constrained polyunsaturated species. Fluorescent avocadyne derivatives were efficiently internalized with distinct localization patterns in L4 larvae and embryonic cells.

**Conclusions:** Overall, the lipid homeostasis remodeling of L4 larvae in response to lipotoxic shock was associated with phospholipid increase and remarkable lipid droplets onset, whereas embryos showed accumulation of lipids in enlarged droplets and developmental arrest.

## Background

Natural compounds are a valuable reservoir of bioactive molecules with significant therapeutic potential. Among these, avocadene and avocadyne, the 17-carbon-long linear fatty alcohols occurring in avocadoes (*Persea americana*) and their acetate derivatives (Fig. 1), have attracted attention due to their unique biological activities [1–9]. Notably, whereas avocadene bears a single terminal unsaturation at the opposite extremity to the alcoholic functions, avocadyne is correspondingly characterized by the presence of a triple bond, which contributes to its distinct biological properties [7, 8] and chemical exploitability. Avocadoes themselves are well recognized for their rich lipid content, vitamins, and antioxidants, which have been related to a range of health benefits [10–13].

**Figure 1.**
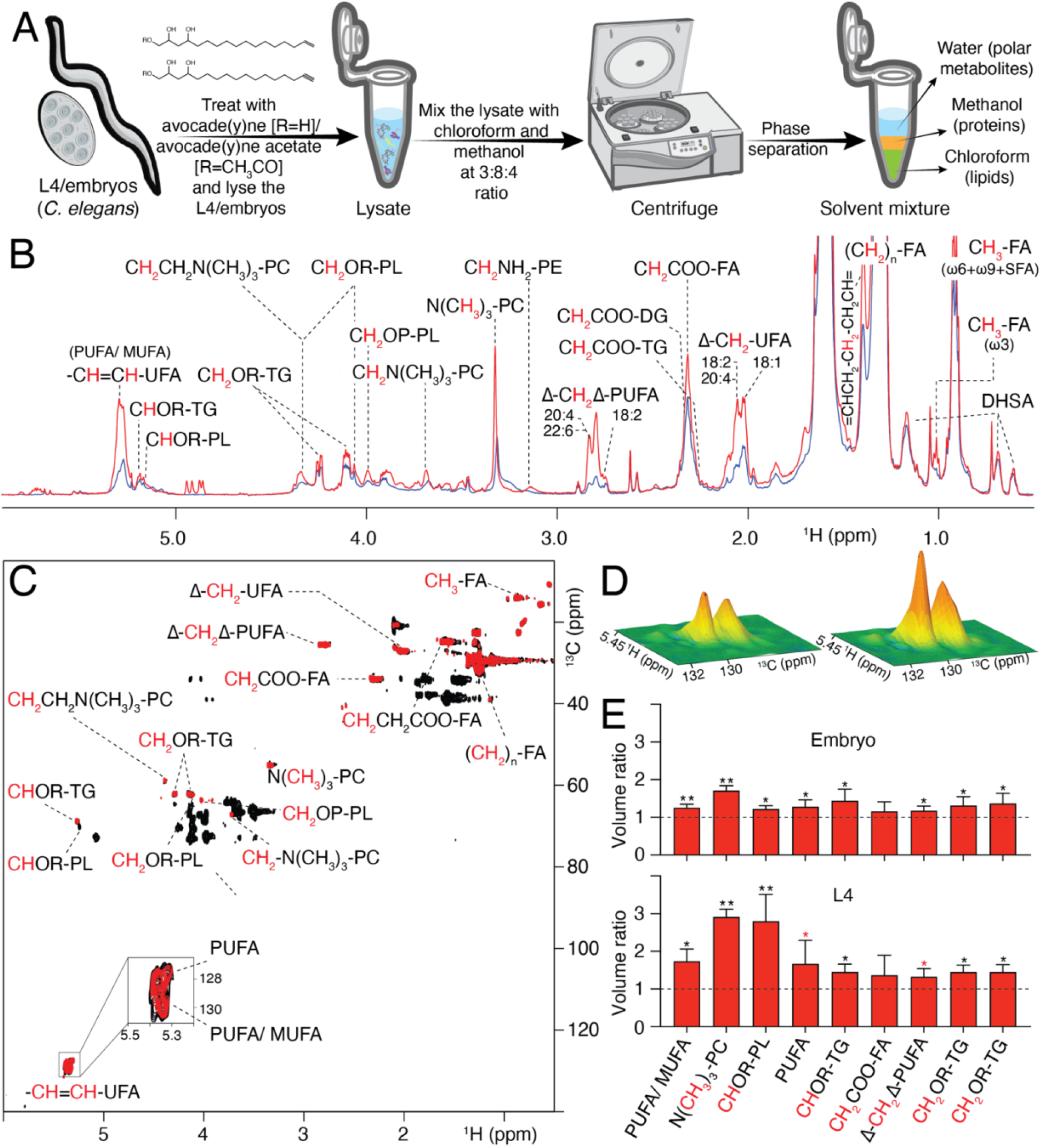
NMR-based lipidomic workflow and quantitative comparison of lipid profiles in control and avocadene-acetate-treated C. elegans L4 and embryos. (A) General workflow to perform lipid extraction according to Folch method. (B) Overlay of 1D ^1^H NMR spectra from control (blue) and treated (red) L4 larvae samples with lipid peak assignments. Contributing protons are marked in red. Abbreviations: FA, fatty acids; UFA, unsaturated fatty acids; MUFA, monounsaturated fatty acids; PUFA, polyunsaturated fatty acids; TG, triglycerides; PL, phospholipids; PC, phosphatidylcholines; PE, phosphatidylethanolamines; DG, 1,2 or 1,3 diacyl-glycerol; DHSA, dihydrosterculic acid; SFA, saturated fatty acid; Δ, unsaturation; ω3, ω6, ω9 = unsaturation position with respect to terminal methyl. (C) 2D ^13^C–^1^H HSQC spectra of L4 larvae lipid extracts; control (red) and treated (black). Peak assignments are indicated, with the contributing proton(s) and carbon(s) highlighted in red. (D) 3D representation of the^13^C–^1^H HSQC cross-peaks of the unsaturated fatty acids (CH=CH-UFA) showing the intensity difference between control (left) and treated (right) L4 larvae samples. (E) Bar graph of cross-peak intensity ratios (treated/control) for the salient lipid signals. The ratio values represent means from three independent biological replicates with errors corresponding to the relative standard deviations. Statistically significant differences are indicated (**=p < 0.01, *=p < 0.05, *=p < 0.08, one-tailed t-test). The denominations PUFA/MUFA and PUFA refer to the olefinic groups (CH=CH-UFA) of the two resolved peaks shown in the inset of panel C and D.

To investigate the biological relevance of these lipids, *Caenorhabditis elegans* (*C. elegans*) provides a highly suitable model system. This nematode is widely used in anti-parasitic drug research because of its genetic tractability, conserved biological pathways with parasitic nematodes, short life cycle, and transparent body which facilitates imaging and molecular studies. In our earlier work, we demonstrated that avocado-derived lipids exert potent nematocidal activity against several parasitic nematodes, including *Brugia pahangi*, *Teladorsagia circumcincta*, *Heligmosomoides polygyrus*, and a multidrug-resistant strain of *Haemonchus contortus* [8]. In *C. elegans*, avocadene, avocadyne, and their acetate derivatives caused concentration-dependent toxicity across different developmental stages, impairing egg hatching, stage 1 development (L1) larval growth, and adult survival at lethal doses, whereas a transient paralysis was observed with stage 4 development larvae (L4) at sublethal doses [8]. Phenotypically, treated worms exhibited fragmented, spherical mitochondria instead of the normal tubular network, consistent with impaired mitochondrial respiration, reduced oxygen consumption, and elevated reactive oxygen species (ROS). These effects suggest that avocado lipids toxicity affects mitochondrial function, along with lipid metabolism and is linked to the previously reported inhibition of acetyl-CoA carboxylase (ACC/POD-2), the rate-limiting enzyme of fatty acid biosynthesis [8, 9].

Building on these findings, the present study aimed to determine how avocado-derived lipids alter the lipid composition of *C. elegans* embryos and L4 larvae, and to explore the interaction of avocado lipids with endogenous lipids. To achieve this, L4 larvae and embryos were treated with avocadene acetate, followed by lipid extraction using the Folch method [14, 15] for detailed lipidomic analysis. In parallel, we employed click chemistry to fluorescently label avocadyne alcohol and acetate (1:1 mixture) with 3-azido-7-hydroxycoumarin, enabling visualization and assessment of avocadyne distribution and interactions.

Our combined strategy, based on *in vitro* lipidomics coupled with thin-layer chromatography (TLC), and *in vivo* fluorescence imaging of labeled derivatives gave complementary information about the molecular interactions and biodistribution of avocadyne. NMR lipidomic analysis revealed increased lipid signals. TLC and NMR further suggested that avocadyne preferentially associates with monounsaturated and flexible polyunsaturated chains of triglycerides, whereas no interactions could be detected with conformationally-constrained polyunsaturated species. Overall, the remodeling of lipid homeostasis following avocadene/yne treatment and lipotoxic shock thereof led to changes in phospholipid levels and lipid droplet profile, with fluorescently labeled subpopulations containing exclusively avocadyne or neutral lipids in larvae, whereas embryos showed enhanced accumulation of neutral lipids in enlarged droplets and developmental arrest. This integrated approach provides insights into the biological roles of avocado lipids in nematodes, and highlights the general utility of combining click chemistry with lipidomics to explore natural compound function.

## Methods

### Chemicals

3-Azido-7-hydroxycoumarin, tetrakis(acetonitrile)copper(I) tetrafluoroborate (Cu(MeCN)₄BF₄), thin layer chromatography (TLC) plates, ethanol, methanol, chloroform, acetic acid (AcOH), hexane, ethyl acetate, trimethylamine and dimethylsulfoxide (DMSO) were purchased from Sigma-Aldrich. Sodium hydroxide, potassium dihydrogen phosphate (KH₂PO₄), disodium hydrogen phosphate (Na₂HPO₄), sodium chloride and hypochlorite (NaClO), and magnesium sulfate (MgSO₄) were also obtained from Sigma-Aldrich. Avocadene acetate was sourced from Microsource Discovery Systems and Sigma-Aldrich. Avocadyne and avocadyne acetate were extracted and purified in-house, as described in our previous publication [7].

### Growing synchronized L4 and embryos

The *C. elegans* N2 (Bristol) strain was used in this study. To obtain synchronized *C. elegans* L4 larvae, gravid adults were treated with a bleaching solution (1% hypochlorite, 0.25 M NaOH) to dissolve adult cuticles and release embryos. Embryos were washed three times in M9 buffer (22 mM KH_2_PO_4_, 42 mM Na_2_HPO_4_, 86 mM NaCl, 1 mM MgSO_4_) and incubated overnight at 20°C with gentle agitation to allow hatching. Synchronized L1 larvae were transferred to nematode growth medium (NGM) plates seeded with *E. coli* OP50 as a food source and cultured at 20°C. Larvae were monitored until they reached the L4 stage (∼48 hours post-hatching). Stage synchronization was confirmed by assessing vulval morphology under a dissecting microscope (40-60x magnification).

For embryo collection, gravid adults were washed from plates using M9 buffer and treated with the alkaline hypochlorite solution (1% NaClO, 0.25 M NaOH) for 5 minutes to dissolve the adult cuticle while preserving embryos that were then pelleted by centrifugation (1,500 × g, 1 minute), washed three times with M9 buffer, and resuspended in fresh M9. Embryo viability and developmental stage were verified using differential interference contrast (DIC) microscopy.

### Treatment and lipid extraction from L4 and embryos

Samples of 10,000 L4 stage worms or 50,000 embryos were exposed for two hours, respectively, to 10 or 20 μM avocadene acetate added as µL aliquots of concentrated solutions in DMSO. Equivalent populations treated with DMSO only were also predisposed as controls. Following the treatment, samples were washed with M9 buffer to remove any residual avocadene acetate and subsequently centrifuged to obtain a pellet of L4 larvae or embryos. Analogous procedures were also used for control samples. The pellet was resuspended in 100 μL of H_2_O and subjected to sonication in vials floating in water/ice bath. The ultrasound treatment was applied by direct immersion of the transducer of a Qsonica Q125-220 sonicator in the sample for three cycles of 15 seconds on and 30 seconds off (frequency 20 kHz, power 25 Watts).

Metabolite extraction was carried out using the Folch extraction protocol [14, 15]. The obtained lysates, kept on ice bath, were then dispersed in a CHCl_3_/CH_3_OH/H_2_O mixture, in volumetric ratio 8:4:3 and final total volume of 2 mL, including the initial aqueous lysate. The samples were vortexed vigorously for 30 seconds four times to favor extraction and ensure homogeneity. The samples were then submitted to a second sonication cycle in an ultrasonic bath pre-filled with ice and water at 40 kHz for 15 minutes. This sonication step enhanced homogeneity and removed debris from the inner tube walls. After sonication, the vials were centrifuged at 4°C for 15 minutes at 1,258 x g. This centrifugation resulted in the formation of three distinct phases: a lower hydrophobic CHCl_3_ phase containing lipids, a middle phase of protein precipitate in a CH_3_OH layer, and an upper hydrophilic phase in H_2_O/CH_3_OH containing water-soluble metabolites.

The supernatant of the lower lipid-containing phase, collected with a Pasteur pipette, was carefully transferred to a new vial and dried under a nitrogen stream. The dried lipid extract was redissolved in 500 µL of chloroform-D (CDCl_3_) for NMR analysis.

### Azide-alkyne cycloaddition reaction

A few µL of concentrated avocadyne and its acetate (1:1) in chloroform was mixed to reach 100 µM with 35 µL of the click reaction mixture, which included 120 µM of 3-azido-7-hydroxycoumarin in ethanol and 2 mM of tetrakis(acetonitrile)copper(I) tetrafluoroborate (Cu(CH_3_CN)_4_BF_4_) in acetonitrile. This mixture was transferred to a 1.5 mL Eppendorf tube and placed on a heating block at 42°C until all the solvent had evaporated and condensed under the tube cap, typically taking about 5 hours. This evaporation step is crucial to ensure the quantitative completion of the click reaction. Subsequently, the Eppendorf tube was centrifuged, vortexed to resuspend any components adhering to the walls, and centrifuged again. The sample was dried and then dissolved in a mixture of water and ethyl acetate to remove copper via phase separation. The ethyl acetate fraction was subsequently dried under nitrogen steam and stored at −80°C.

### Lipid extraction and click reaction

A population of 20,000 L4 worms was initially exposed to 12.5 μM of avocadyne acetate for one hour. This concentration was then increased to 20 μM, and the treatment continued for an additional four hours. For embryos, 100,000 individuals were exposed to 20 μM of avocadyne acetate for two hours. Following the treatment, the worms or embryos were washed with M9 buffer to remove any residual avocadyne acetate and subsequently centrifuged to obtain a worm or embryo pellet for the Folch extraction already described. A 5 µL aliquot of this lipid extract was combined with 35 µL of the click reaction mixture to undergo the clicked-avocadyne synthesis procedure above mentioned. The product was applied onto a 20 cm × 20 cm silica TLC plate. The plate was initially developed using a solvent mixture system composed by CHCl_3_/CH_3_OH/water/AcOH in a ratio of 68.4/26.3/4.2/1.1, allowing the solvent to travel 8 cm above the plate-mixture meniscus. The plate was then dried by flushing nitrogen gas and developed a second time using a 1:1 mixture of hexane and ethyl acetate and allowing the solvent front to travel 18 cm. After the second development, the plate was dried again with nitrogen gas. To neutralize any acidic effects, the plate was soaked in 4% trimethylamine in hexane. Finally, the plate was dried under a hood by nitrogen steam to remove any remaining solvent. The dried plate was then used for fluorescence imaging, with excitation and emission wavelengths at 350 and 480 nm, respectively.

### Fluorescence imaging of worms and embryos

*C. elegans* L4 larvae or embryos were incubated with avocadyne conjugated to 7-hydroxycoumarin azide for 30 minutes at room temperature. Following treatment, worms or embryos were washed three times with M9 buffer to remove unbound compounds. A subset of worms or embryos was then spread onto a 2% agarose pad mounted on microscope glass slide for imaging. Immobilization was performed by dropping 10 mM levamisole to minimize movement during image acquisition. Fluorescence imaging was carried out using a Leica DMi8 inverted fluorescence microscope equipped with a DAPI filter set (excitation: 350/405 nm, emission: 480 nm), suitable for detecting fluorescence from avocadyne conjugated to 7-hydroxycoumarin azide. Images were captured using a Leica DFC camera and processed using Leica Application Suite X (LAS X) software. Exposure times and gain settings were kept constant across samples to allow for direct comparison.

### BODIPY™ 493/503 staining and fluorescence imaging

To observe the lipid accumulation in the adult germline, young adults were treated with DMSO or 25µM avocadene acetate for 1 hour, followed by staining in BODIPY™ 493/503 solution for 30 minutes. After M9 washes, fluorescent images were captured using a Leica DMi8 microscope with 60X oil objective. The transgenic strain *VS50 plin-1(hj178[PLIN-1::GFP*]), kindly gifted by Prof Ho Yi Mak, was used to visualize the lipid droplets and to assess the incorporation of conjugated avocadyne in embryos as PLIN-1-GFP (perilipin–green fluorescent protein) is a fluorescent marker to outline the LD surface [16]. The embryos were dissected from young adults and treated with DMSO or 50µM avocadene acetate, and with 50-75 µM 3-azido-7-hydroxy coumarin or hydroxycoumarin-tagged avocadyne for 30 minutes. The embryos were then transferred to a 2% agarose pad and microscopy analysis was performed using Andor Dragonfly 600 Spinning Disk Confocal microscope with 100X oil objective (excitation: 488 nm).

### Live imaging of lipid droplets by fluorescence microscopy

Synchronized L1-stage worms with DHS-3-GFP (short-chain Dehydrogenase/reductase–green fluorescent protein) fluorescently tagged intestinal lipid droplets *[LIU1(dhs-3p::DHS-3::GFP)]*, provided by Caenorhabditis Genetics Center] were cultured in 96-well flat-bottom plates in a total volume of 100 µl per well, consisting of 70 µl M9 buffer and 30 µl OP50 bacterial suspension. Worms were allowed to develop to the L4 stage. Hydroxycoumarin-tagged avocadyne (treatment) or 3-azido-7-hydroxycoumarin (control) in DMSO, or just DMSO was then added to a final concentration of 20 µM and incubated for 1 hour. Following incubation, worms were mounted on 3% agarose pads on glass slides and immobilized with 2–3 µl of 1 mg/mL levamisole in M9 buffer. Fluorescence images were acquired using a Leica DMi8 microscope with excitation at 488 nm (for GFP) and 405 nm (for hydroxycoumarin-tagged avocadyne). Images were processed with ImageJ.

### NMR spectroscopy

All the samples (avocadyne, avocadyne acetate, their hydroxycoumarin-linked conjugates and/or TLC extracts) were dissolved in 550 µL CDCl_3_ and transferred into 5 mm NMR tubes. NMR experiments were performed using a Bruker Avance III spectrometer operating at a magnetic field strength of 14.0 T (¹H Larmor frequency of 600.19 MHz) and equipped with a triple-resonance cryoprobe. One-dimensional ¹H NMR spectra were acquired using the first *t_1_* increment of a standard NOESY [17] sequence. Free induction decays (FIDs) were recorded over a spectral width of 16 ppm, 32768 data points, 256 transients, and a relaxation delay of 1, 2 or 10 s. FIDs were processed using a squared cosine bell or an exponential multiplication weighing function with 0.3 Hz line-broadening factor prior to Fourier transformation.

2D ¹H-¹H total correlation experiments (¹H-¹H TOCSY) were collected to confirm the lipid assignments [18–22]. The spectra were acquired using the DIPSI2 sequence for homonuclear Hartmann-Hahn transfer, in the phase-sensitive mode and without water suppression [23]. Data matrices of 1024/2048 × 128/256 points (*t_2_* x *t_1_*) were recorded, with 64 transients per FID, 1.0 s relaxation delay and 60-100 ms TOCSY mixing time.

2D ^13^C-¹H Heteronuclear Single Quantum Coherence (^13^C-¹H HSQC) spectra [24] were acquired using the hsqcedetgpsisp2.3 pulse sequence of the Bruker library. The experiment was performed with sensitivity enhancement and phase-sensitive detection using Echo/Antiecho-TPPI gradient selection [25, 26]. Double INEPT transfer with trim pulses was employed, and multiplicity editing was applied during the selection step, with suppression of long-range coupling connectivities. Matched sweep adiabatic pulses were used for all 180° pulses on the X-nucleus channel, and gradients were incorporated in the back-INEPT transfer [27–29]. ^13^C-¹H HSQC spectra were also acquired without multiplicity editing during the coherence selection step (hsqcetgpprsisp2.2 pulse sequence). Data sets of 1024 × 256 points were collected, with 256 transients per FID and 1.0 s relaxation delay. For the clicked-lipid extracts, spectra were acquired over 1024 × 128 matrices, with 448 transients per FID and the same relaxation delay.

### Mass spectrometry

The samples suspended in 50% aqueous acetonitrile (v/v) were analyzed by nanoMate-coupled Fourier transform ion cyclotron resonance mass spectrometry (FT-ICR-MS). The nanoMate is an automated, chip-based electrospray ionization (ESI) source that provides highly stable, low-flow nanospray, improving ionization efficiency and run-to-run reproducibility while minimizing cross-contamination between samples. Sample volume of 10 µl was injected with a spray voltage set of 1.7 kV and gas pressure 0.2 psi. FT-ICR-MS enables high-resolution, accurate-mass measurement and isotopic fine structure analysis, facilitating unambiguous compound characterization. The high-resolution mass spectra were acquired on a Bruker solariX FT-ICR mass spectrometer (7.0 T magnet, Bruker Daltonics, Germany) in positive ion mode over the *m/z* range 100–1800 Th, with 100 scans (0.1 sec per scan) accumulated per sample. Data were acquired using ftmsControl and processed with Bruker DataAnalysis software.

### Toxicity assay on *C. elegans*

Synchronized embryos were used to get first larval stage (L1) worms. Dose-response experiments were then conducted by dispensing 20 μL of M9 containing 10-15 L1 larvae and 30 μL of OP50 into each well of a 96-well plate. This was followed by the addition of equimolar mixtures of avocadyne and avocadyne acetate or their hydroxycoumarin conjugated from 0 to 100 μM (total concentration). The tested intermediate concentrations used were 1.25, 2.5, 5, 10, 12, and 50 μM. The growth of the worms was monitored until they reached adulthood and gave rise to F1 progeny. The experiments were conducted in triplicate, calculating the average and standard deviation for each assay. The number of L1 larvae that developed to adulthood was plotted against the logarithm of the total concentration of the tested compounds. Nonlinear regression analysis was performed with GraphPad Prism 10 using a sigmoidal dose-response model to fit the data and calculate the LD_50_ (median lethal dose) values and relative errors (92% confidence interval).

## Results

### Avocadene acetate treatment induces lipid accumulation in *C. elegans* larvae and embryos

To investigate the lipid profile alterations associated with avocadene acetate-induced transient paralysis and mitochondrial damage of *C. elegans* L4 larvae and embryonic growth arrest [8], control and avocadene-acetate-treated (10-20 µM, 8-2 h) L4 larvae and embryos underwent lipid extraction by the three-solvent Folch procedure (Fig.1A). The NMR spectra of the resulting lipid extracts were collected in CDCl_3_ for detailed structural and composition analysis. Overall, the NMR analysis revealed a consistent and notable increase of lipid signal intensity in the treated samples compared to the controls, particularly with L4 larvae The ^1^H NMR trace overlay of Fig. 1B clearly shows the increased lipid content in the treated L4 larvae extract. A similar pattern was observed also for the embryos extract (Fig. S1B). The ^1^H peak assignments were obtained from previously reported results [18–22]. The typical NMR spectrum profile shows signals corresponding to various lipid types, including polyunsaturated and monounsaturated fatty acids (PUFA/MUFA), whose peaks are sometimes unresolved as observed with the characteristic olefinic signals, i.e. those from the -CH=CH-groups, collectively referred to as unsaturated fatty acids (UFA). Additional poorly resolved signals arise from the CH_2_ in position 2 (or a) of saturated fatty acids (SFA) and UFA with unsaturation far from the free or esterified carboxyl group. On the other hand, we distinctly observe the characteristic trimethylammonium peak from the phosphocholine headgroup of phosphatidylcholines (PC), i.e. N(CH₃)₃-PC, along with proton signals from the central (CHOR) and external (CH_2_OR) glycerol backbone of glycerophospholipids and triacylglycerols, shortly named phospholipids (PL) and triglycerides (TG), respectively. Another characteristic pattern arises from the allylic methylene hydrogens (Δ-CH_2_ with Δ standing for unsaturation) of MUFA and PUFA with large separations (≥ 2 C atoms) between double bonds, and the bis-allylic methylene protons of polyunsaturated fatty acids with single-carbon spacing between unsaturations (Δ-CH₂-Δ). Further details are reported in Fig. 1 and relative caption.

To improve the analysis resolution, we collected 2D ^13^C-^1^H HSQC spectra. Based on three-replicate experiments on independent samples, 2D integration of ^13^C-^1^H HSQC spectra and subsequent evaluation of the Intensity ratios between corresponding peaks from treated and control samples quantitatively established, within the limits of extraction variability and signal overlap, the higher lipid content levels in the treated L4 larvae (Fig. 1C, D, E). Analogous, albeit reduced, enhancements were also assessed by 2D integration of embryos spectra (Fig. S1C, D, Fig. 1E). In particular, an appreciable increase of TG intensity could be measured from the treated larvae spectra (Fig.1C, D), as readily appreciated also in the corresponding 1D trace overlay (Fig.1B), and the treated embryo spectra (Fig. S1B, C, D). In addition, we also detected much more significant increases of UFA levels in treated larvae (Fig. 1B, C, D), but only slight increments in treated embryos (Fig. S1B, C), according to the UFA allylic CH_2_ peak of mono and polyunsaturated species (18:1, 18:2, 18:3, 20:4, etc.) and PUFA bis-allylic peak (Δ-CH₂-Δ in 18:2, 18:3, 20:4, 22:6, etc.), suggesting altered fatty acid catabolism and anabolism. Consistently with the trend of most lipid species moieties, the saturated and unsaturated fatty acid αCH_2_ and terminal CH_3_ signals also showed higher intensities, though no specific quantitative inference could be drawn because of the unresolvable merging of contributions from many classes of chains. Instead, based on the variations of signature-signals, L4 larvae exhibited relevant increments of PL and less pronounced increments of PUFA and MUFA chains of TG or PL (Fig. 1E). Characteristic PL signals also showed slightly increased abundance of the species in treated embryos. Finally, both L4 larvae and embryos showed a distinctly large and meaningful increase of the phosphocholine moiety signal from PC (Fig. 1E). The increment of PL and PC levels, however, was much more substantial in L4 larvae compared to embryos (Fig. 1E), and suggests changes in membrane lipid composition or turnover. Overall, this pattern implies remodeling or dysregulation of lipid handling and mobilization.

These results are consistent with our previous findings on mitochondrial damage and acetyl-CoA carboxylase (ACC/POD-2) inhibition upon treatment with avocadene/yne and their acetate derivatives [8, 9] Acetyl-CoA carboxylase is the key enzyme responsible for the formation of malonyl-CoA, which, in turn, is the first step of fatty acid biosynthesis and simultaneously regulates fatty acid catabolism by inhibiting their transport in mitochondria by carnitine palmitoyl transferase I (CPT-I) [30–32].

At first glance, therefore, the inhibition of an enzyme expected to arrest anabolism and activate mitochondrial catabolism of lipid fatty acids [33, 34] might seem inconsistent with the lipid increments observed by NMR. However, ACC inhibition can produce paradoxical lipid redistribution effects due to compensatory remodeling of lipid levels [35–37]. The paradoxical lipid increments were attributed to the activation of anaplerotic pathways by increased levels of acetyl-CoA and tricarboxylic acid cycle intermediates [37]. The partial rescue of the treated L4 larvae by supplementation with malonyl-CoA or the antioxidant N-acetylcysteine, or by knockdown of the carnitine shuttle genes using RNA interference [8], demonstrates that the primary toxic effect of ACC/POD-2 inhibition by avocadene/yne compounds involves mitochondrial damage including impaired respiration, reduced oxygen consumption, and elevated levels of reactive oxygen species (ROS) [8]. Lipid catabolites also accumulate as a consequence of the ensuing mitochondrial dysfunction, rather than being cleared through efficient β-oxidation as expected from the lack of malonyl-CoA inhibition of CPT-I [30–32]. The detergent-like properties of the accumulated species [38–41] affect neuromuscular function and thus contribute to the onset of transient paralysis in L4 larvae. The animals, however, counteract the disruption of lipid homeostasis and consequent paralysis by triggering enhanced lipid storage mechanisms to mitigate lipotoxic effects, as examined below.

### Avocadene acetate – endogenous lipid interaction

The alteration of lipid homeostasis inferred from the NMR analysis also showed an interesting feature concerning the interaction of avocadene acetate with the endogenous lipid pool of *C. elegans* L4 larvae and embryos.

In our earlier study [8], we reported the presence of avocadene acetate in the chloroform phase of the Folch extracts and highlighted the similar values (within the experimental error) of the diffusion coefficients (*D*) for the initial and final positions of the avocadene acetate molecule, according to NMR DOSY measurements [42, 43] that were carried out on solutions of the pure compound and the extracts from treated *C. elegans* larvae and embryos [8]. This evidence is consistent with the molecular integrity of the incorporated avocadene acetate, i.e. absence of metabolic transformation. However, no interpretation was provided for the actual *D* values that were experimentally determined for the pure compound and the extract solutions. On considering those earlier reported values, a distinct, yet consistent, reduction emerges for the diffusion parameters in chloroform solution of the biological extracts compared to the pure compound (Table 1).

**Table 1.**
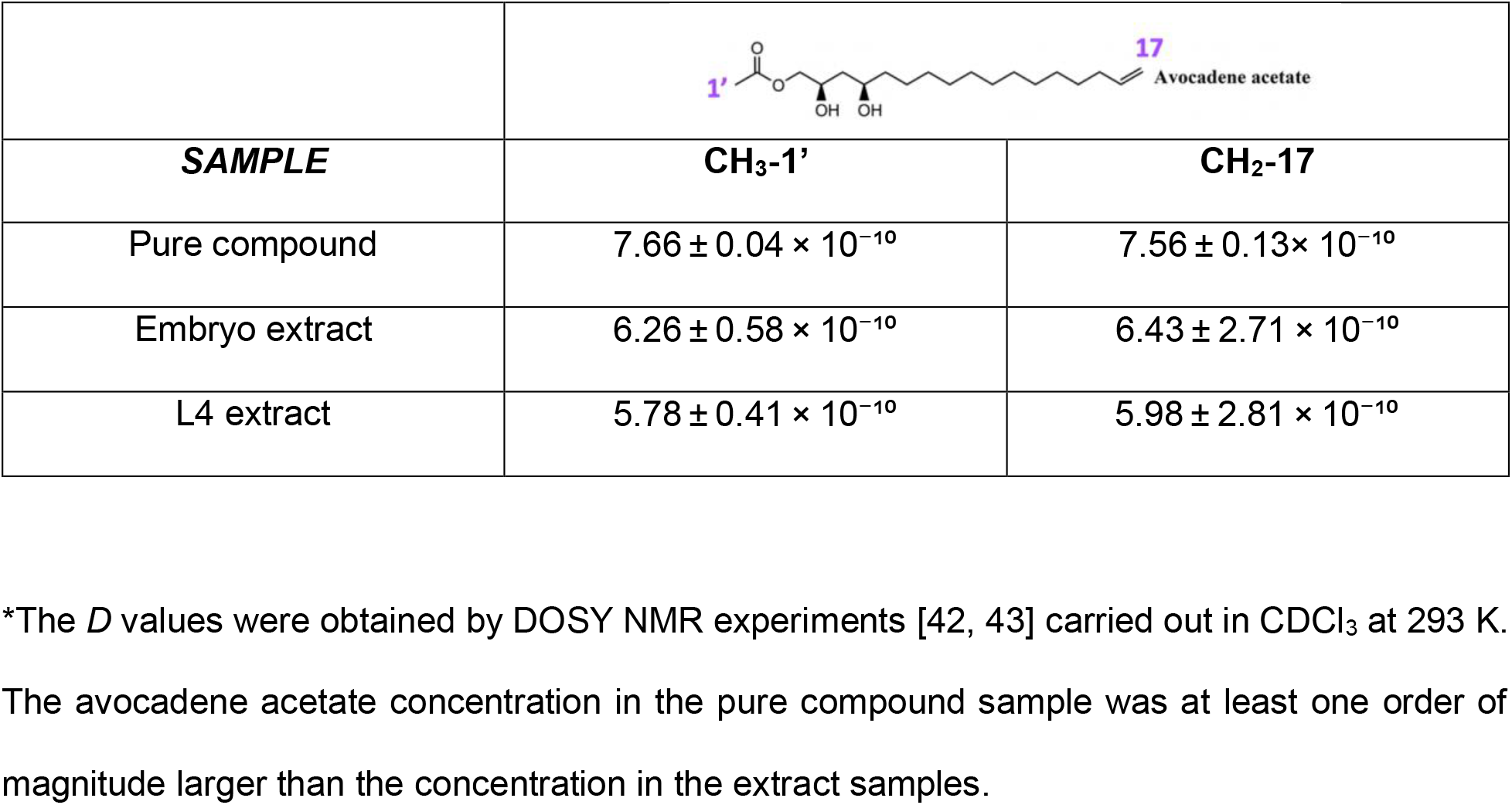
Diffusion coefficient (*D*) values (in m²/s) measured for the extreme molecular locations of avocadene acetate*.

**Table 1.**
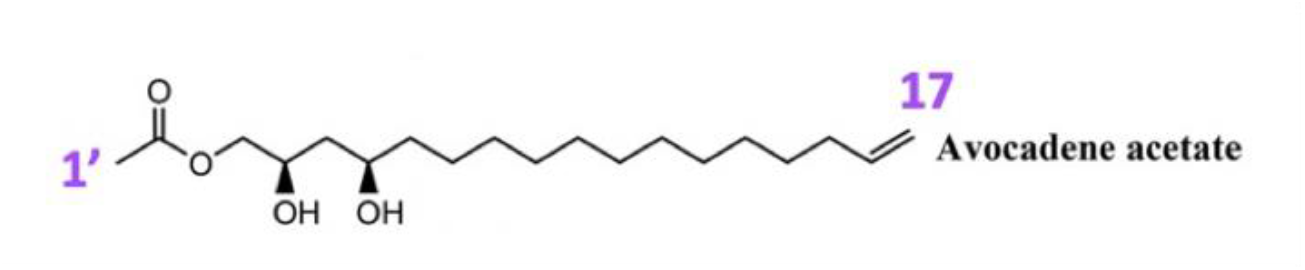
Scheme.

The measurable decrease in diffusion rates and the substantial invariance of the chemical shifts suggest that avocadene acetate dynamically associates with other components, essentially lipidic, that are present in the extracts. These observations provide evidence supporting the possible occurrence of interactions with endogenous lipids also in the biological *milieu*.

### Conjugation of avocadyne with 3-azido-7-hydroxycoumarin

7-Hydroxycoumarin, also known as umbelliferone, skimmetine, and hydrangine, is a natural compound belonging to the coumarin family, characterized by the precursor chromophore fluorescence. However, the fluorescence of 7-hydroxycoumarin is quenched when an azide functional group is introduced in position 3 of the coumarin scaffold, forming 3-azido-7-hydroxycoumarin. This quenching effect can be reversed by the copper(I)-catalyzed Huisgen cycloaddition reaction between the azide and an alkyne, resulting in the formation of a 1,2,3-triazole derivative that restores the fluorescence. The reaction occurs quantitatively in a single step and belongs to a class of quick and facile transformations referred to as click chemistry.

This approach has been widely applied in the synthesis of fluorescent dyes due to the high efficiency of the reactions and general applicability for biomolecular labeling. In this study, we optimized the click reaction between avocadyne and 3-azido-7-hydroxycoumarin (Fig. 2A), as described in the Methods section. Successful addition of 3-azido-7-hydroxycoumarin to the positions 16 and 17 of avocadyne was initially confirmed by the strong cyan fluorescence of the resulting product under UV light (365 nm; Fig. 2B). This fluorescence arises from the conjugation of the azido group with formation of the triazole ring, which restores the fluorescent properties of the coumarin scaffold. Solubility tests in methanol, acetonitrile, ethanol, chloroform, and DMSO indicated that the reaction product was fully soluble in all these solvents. To further confirm the structural modifications, we compared the ¹H NMR spectra of the avocadyne reactant and triazole derivative product (Fig. 2C). Notably, the signals at 1.94 and 2.18 ppm corresponding to the protons at positions 15 and 17 of avocadyne, respectively, were absent in the spectrum of the reaction product because of the alkyne moiety quantitative transformation into the addition product. The excitation and emission properties of the fluorescently tagged avocadyne were then determined. The conjugate exhibited distinct excitation and emission maxima (Fig. 2D), as expected for the spectral profile of coumarin derivatives.

**Figure 2.**
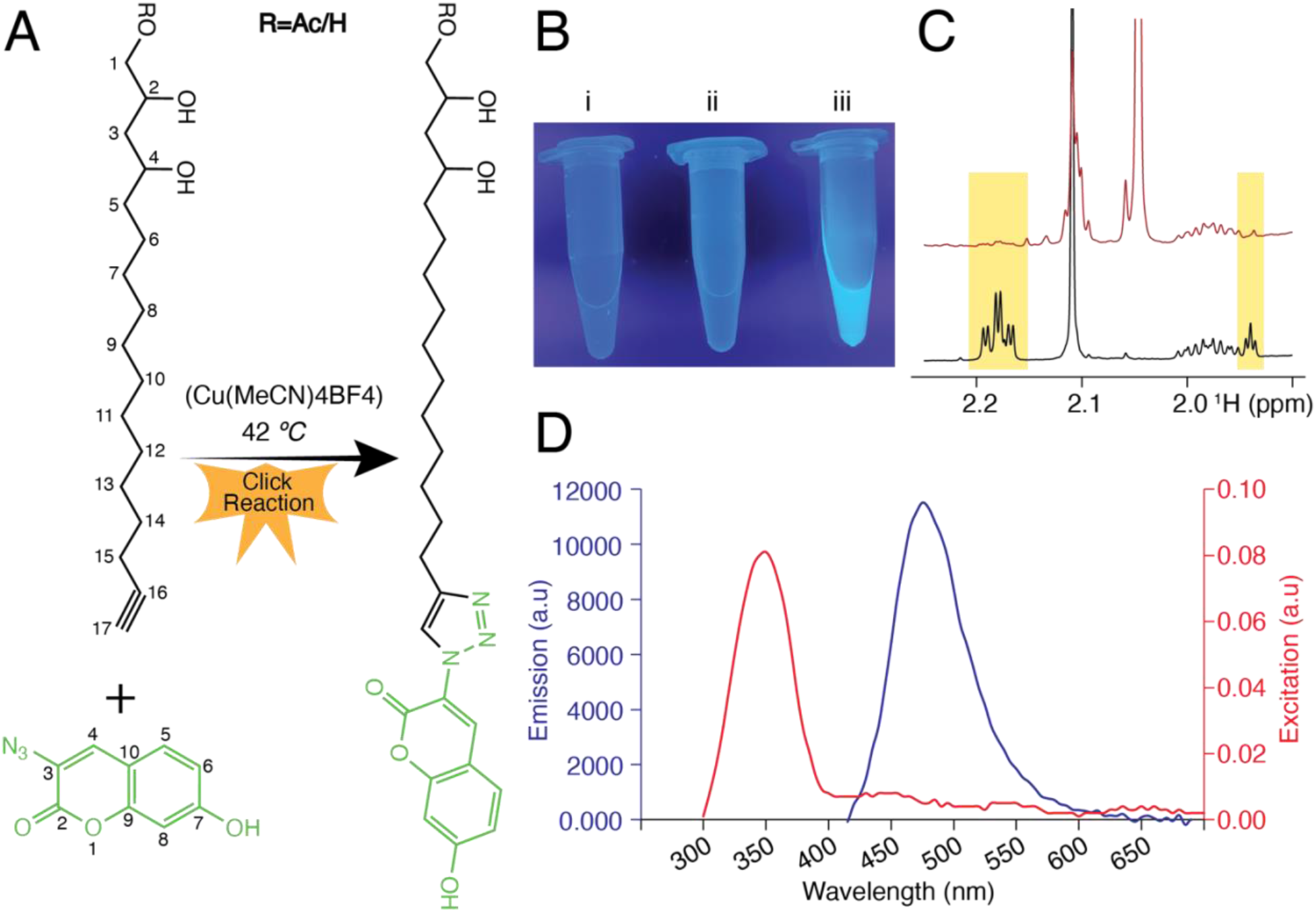
Click-chemistry coupling of avocadyne with 3-azido-7-hydroxycoumarin and product analysis. (A) General diagram of the click-chemistry reaction between avocadyne or its acetate derivative (black) and 3-azido-7-hydroxycoumarin (green) (Ac = acetyl). (B) UV lamp irradiation at 365 nm of the avocadyne-hydroxycoumarin conjugate product (iii) and isolated reactant controls, (i) avocadyne and (ii) 3-azido-7-hydroxycoumarin. (C) comparison of the ^1^H NMR spectra of avocadyne (black) and the avocadyne-hydroxycoumarin conjugate (brown). The yellow shading highlights the signals of avocadyne hydrogens in positions 15 and 17 that disappear in the product (numbering in panel A). (D) Excitation at λ_ex_ = 350 nm (red) and emission at λ_em_ = 480 nm (dark blue) of the avocadyne-hydroxycoumarin conjugate fluorescence spectrum.

### Fluorescence-assisted identification of lipid Interactions with avocadyne and its derivative

Avocadyne, in both free alcohol and acetate derivative forms, exhibits cytotoxic effects on *C. elegans* larvae and embryos as shown through biological and biochemical assays [8]. To investigate the underlying mechanisms, prompted by the diffusion rate reduction of avocadene acetate in embryos and larvae extracts, we further checked whether free avocadyne and its acetate interact with lipids from *C. elegans* L4 larvae and embryos. The chemicals were supplemented at 1:1 ratio and 20 µM to L4 or embryos suspensions for 3-5 hours, followed by Folch extraction to isolate and dry the chloroform phases. The dried lipid extracts were split into small aliquots and reacted with 3-azido-7-hydroxycoumarin in the presence of Cu(I), to conjugate the chromophore to the avocadyne compounds present in the lipid extract for selective fluorescence detection. When inspected under UV light (365 nm), the reaction mixtures exhibited strong cyan fluorescence, consistent with the characteristic emission of coumarin derivatives (Fig. 3A, Fig. S2A).

**Figure 3.**
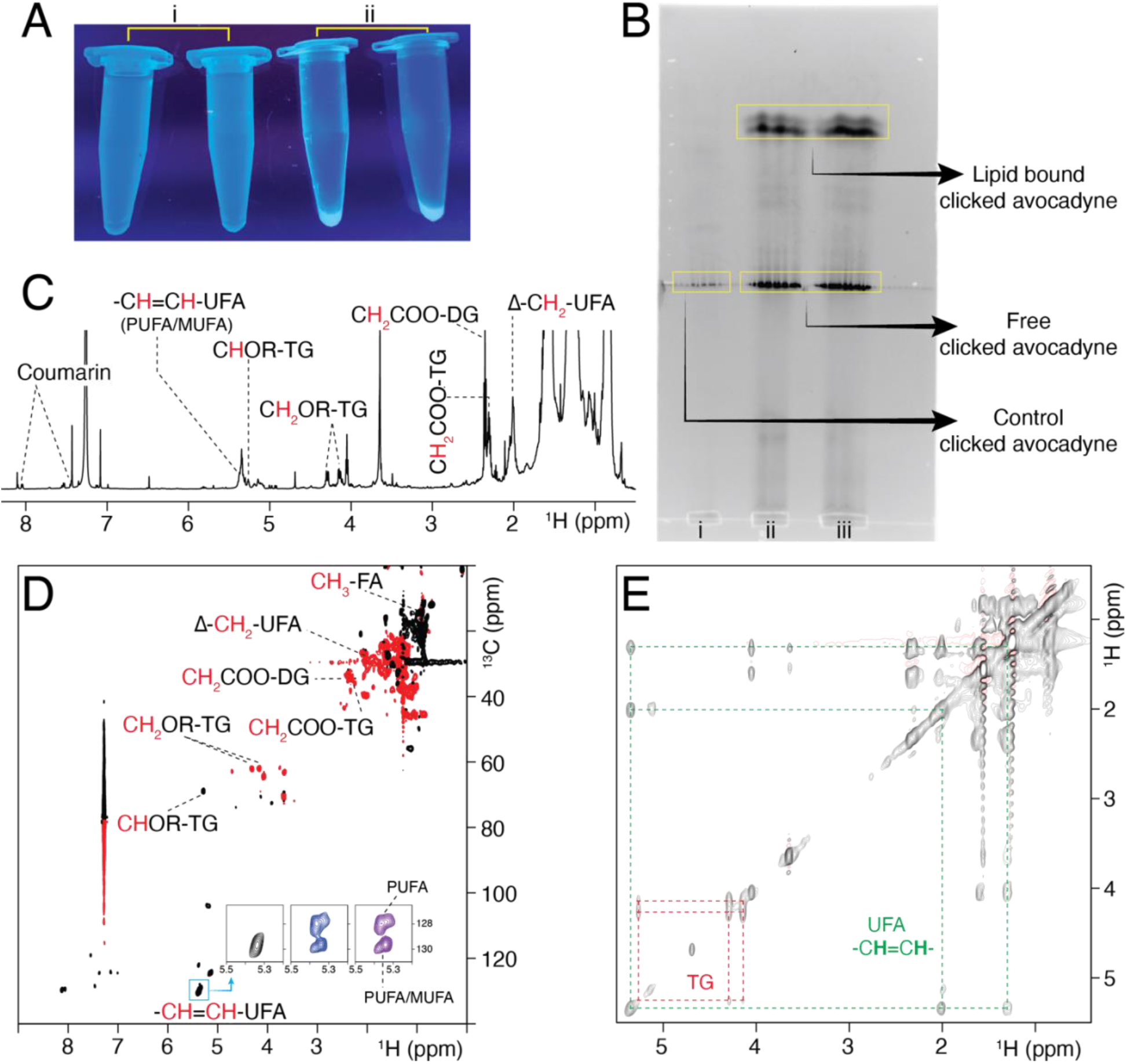
Fluorescence and NMR tracing of avocadyne-treated embryo extract after fluorophore conjugation and TLC separation. (A) UV lamp irradiation (365 nm) of avocadyne-treated C. elegans embryo suspensions before (i) and after (ii) conjugation to the coumarin derivative fluorophore (see Fig. 2). (B) TLC plate development highlighting fluorescent strip separation corresponding to free and lipid-bound hydroxycoumarin-tagged avocadyne. To enhance fluorescence detection, the TLC plates were briefly immersed in hexane containing N,N-di-isopropylethylamine (Hünig’s base), a volatile, non-fluorescent base ensuring that the hydroxycoumarin dye of the conjugated species remains in the highly-fluorescent phenolate form, to significantly amplify the emission intensity. (C)^1^H 1D spectrum in CDCl_3_ of the Folch extract from avocadyne-treated embryos after submission to coumarin derivative conjugation, TLC resolution and chloroform extraction selectively carried out on fluorescent TLC bands. The lipid assignments are reported. (D) ^13^C-^1^H edited HSQC spectrum showing the assignments of the relevant lipid signals. The insets indicate the resonances corresponding to the UFA olefinic nuclei (CH=CH-UFA) that were observed in different samples, namely lipids extracted from fluorescent TLC bands (black), lipids from untreated embryos (blue), and lipids from avocadyne-treated embryos (purple). (E) ^1^H-^1^H TOCSY spectrum highlighting triglycerides (TG) and unsaturated fatty acids (UFA) connectivities. The corresponding results obtained with C. elegans L4 larvae are reported in Fig. S2.

Subsequently, the reaction mixtures were chromatographically resolved on standard silica gel TLC plates using a dual-solvent system of varying polarity. As a small and relatively hydrophobic fluorophore, 7-hydroxycoumarin could enable efficient TLC separation of the labeled components and their lipid-association adducts with minimal effect on their migration behavior.

The fluorescence of a typical TLC plate is documented in Fig. 3B (and Fig. S2B). The silica from the fluorescent band areas was carefully scraped off and submitted to chloroform extraction. The extracted product was then dried, redissolved in CDCl_3_ and analyzed by NMR (Fig. 3C, Fig. S2C). The spectrum revealed characteristic signals corresponding to lipids and coumarin. In particular, we observed resonances from the olefinic protons of monounsaturated and polyunsaturated fatty acids (CH=CH-UFA), signals from triglycerides’ glycerol moiety (CHOR-TG and CH₂OR-TG), methylenes next to carboxyester groups of acyl chains (CH_2_COO-DG, CH_2_COO-TG) of unsaturated and saturated di- and triglycerides, as well as allylic methylenes (Δ-CH₂) of PUFA and MUFA. These assignments confirm the presence of both saturated and unsaturated chains in neutral lipid species within the silica-bound lipid fraction. Interestingly, we did not observe the characteristic bis-allylic signal (Δ-CH₂Δ-PUFA) at 2.8 ppm, suggesting that avocadyne does not interact with PUFA with double bonds separated by a single methylene. This observation was further supported by the 2D ¹³C-¹H HSQC spectrum, where signals typically associated with that type of PUFA at 127.9 ppm and 5.36 ppm (olefinic region), as well as at 25.85 ppm and 2.8 ppm (bis-allylic methylene region), were absent (Fig.3D, S2D). Instead, we detected clear signals at 129.7 ppm and 5.34 ppm, corresponding to the olefinic protons (CH=CH-UFA) of MUFA and/or PUFA with two or more methylene between double bonds. Consistently, the ¹H-¹H TOCSY spectrum revealed strong correlations between the olefinic protons and both the Δ-CH₂ protons of UFA and the CH₂ envelope of fatty acid chains, but no correlations with the bis-allylic methylene resonances centered at 2.8 ppm (Δ-CH₂-Δ-PUFA) (Fig. 3E, Fig. S2E). Altogether, these results indicate that avocadyne preferentially associates with SFA and with UFA containing either a single double bond or multiple double bonds separated by at least two methylene groups. This selectivity appears to depend on lipid-chain conformational dynamics: the constraints imposed by double bonds separated by a single bis-allylic methylene group are sufficient to reduce the interaction, if present, below the detection threshold.

The NMR datasets of the TLC selective extraction products (¹H, ¹³C-¹H HSQC, and ¹H-¹H TOCSY) were further analyzed with the COLMAR online server, employing a lipid metabolite database obtained in chloroform [44]. This analysis revealed a more diversified profile of lipid species in the fluorescence-labeled fraction, including monounsaturated and saturated fatty acids such as oleic, elaidic, undecanoic, nervonic, decanoic, nonanoic, and palmitoleic acids. In addition, triglycerides were identified, including TG(10:0/10:0/10:0) and TG(16:0/16:0/16:0). These assignments are summarized in Table S1. With the exception of the monoterpene alcohol, the COLMAR profile is compatible with the assignments obtained from established literature evidence. The TLC selective extraction products from embryos and L4 larvae were also submitted to mass spectrometry analysis (Fig. S4, Fig. S5). These controls could detect triglycerides TG(10:0/10:0/10:0) and TG(16:0/16:0/16:0), the free and conjugated avocadyne and avocadyne acetate, along with the free fatty acids, except nonanoic and undecanoic acids. No avocadyne covalent adduct with diglycerides DG(10:0/10:0) and DG(16:0/16:0) was detected.

### Biological activity of avocadyne compounds and their hydroxycoumarin conjugates

A preliminary comparison of the predicted physicochemical properties of avocadyne and avocadyne acetate with those of their corresponding hydroxycoumarin conjugates was performed using the SwissADME algorithm [45]. In addition to the expected differences in molecular size and unsaturation, hydroxycoumarin conjugation was predicted to increase affinity for polar environments, but this effect could possibly be balanced by permeability reduction (Fig. S3). Consistent with these changes, conjugation also reduced biological activity. Our previous study reported LD₅₀ values of 12.2 ± 1.0 µM and 5.2 ± 0.2 µM for L1 developmental arrest by avocadyne and avocadyne acetate, respectively [8]. An equimolar mixture of the two compounds had an LD₅₀ of 10.2 ± 0.9 µM, compared with the value of approximately 7.3 µM predicted under a concentration-addition model. The higher experimentally observed LD₅₀ therefore indicates an antagonistic interaction between the two compounds. The corresponding equimolar mixture of hydroxycoumarin conjugates exhibited an even higher LD₅₀ of 15.7 ± 1.7 µM (Fig. 4), likely reflecting the increased polarity and reduced permeability associated with hydroxycoumarin conjugation, together with the antagonistic behavior of the component compounds.

**Figure 4.**
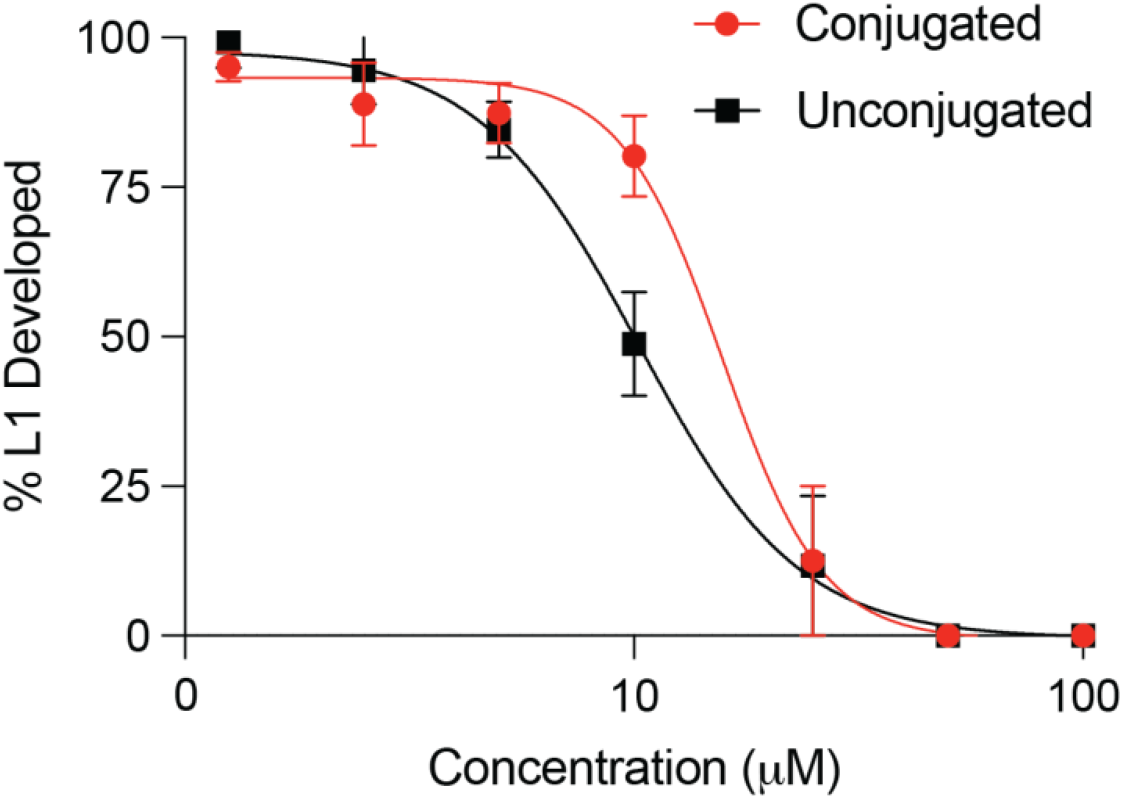
Dose-response curve of C. elegans L1 developmental inhibition. The black curve represents the effect of an equimolar mixture of avocadyne and avocadyne acetate (Unconjugated), while the red curve indicates the result obtained with an equimolar mixture of the corresponding hydroxycoumarin-conjugated species. The error bars are standard deviations of average experimental values (n = 3). The C. elegans N2 (Bristol) strain was used for all assays.

### Localization of tagged avocadyne in *C. elegans* L4 larvae and embryos

To investigate the tissue localization and the relative time progression of avocadyne in *C. elegans*, we monitored the molecule linked to a fluorescent tag (3-azido-7-hydroxycoumarin) and analyzed its distribution within L4 larvae and embryos using fluorescence microscopy. The microscopy images revealed a distinct signal, confirming the internalization of avocadyne. The fluorescence was localized in specific organism regions, suggesting potential interactions with lipid-rich organelles such as lipid droplets (LD) or different tissue compartments.

In L4 larvae, fluorescence from hydroxycoumarin-tagged avocadyne was first detected in the pharyngeal region and then the intestinal lumen within 30 minutes of incubation (Fig. 5A). Diffusion of conjugated avocadyne into embryos outside the intestinal lumen was also observed at a later stage in gravid adult animals (Fig. S6). The prominent fluorescence in the pharynx and intestinal lumen suggests that conjugated avocadyne is ingested during feeding and transiently accumulates as it passes through the digestive tract. This early localization coincides with the previously observed rapid effects of avocadene/yne treatment, including mitochondrial dysfunction and transient paralysis, and is consistent with internalization through the intestine, the principal site of food processing and nutrient absorption.

**Figure 5.**
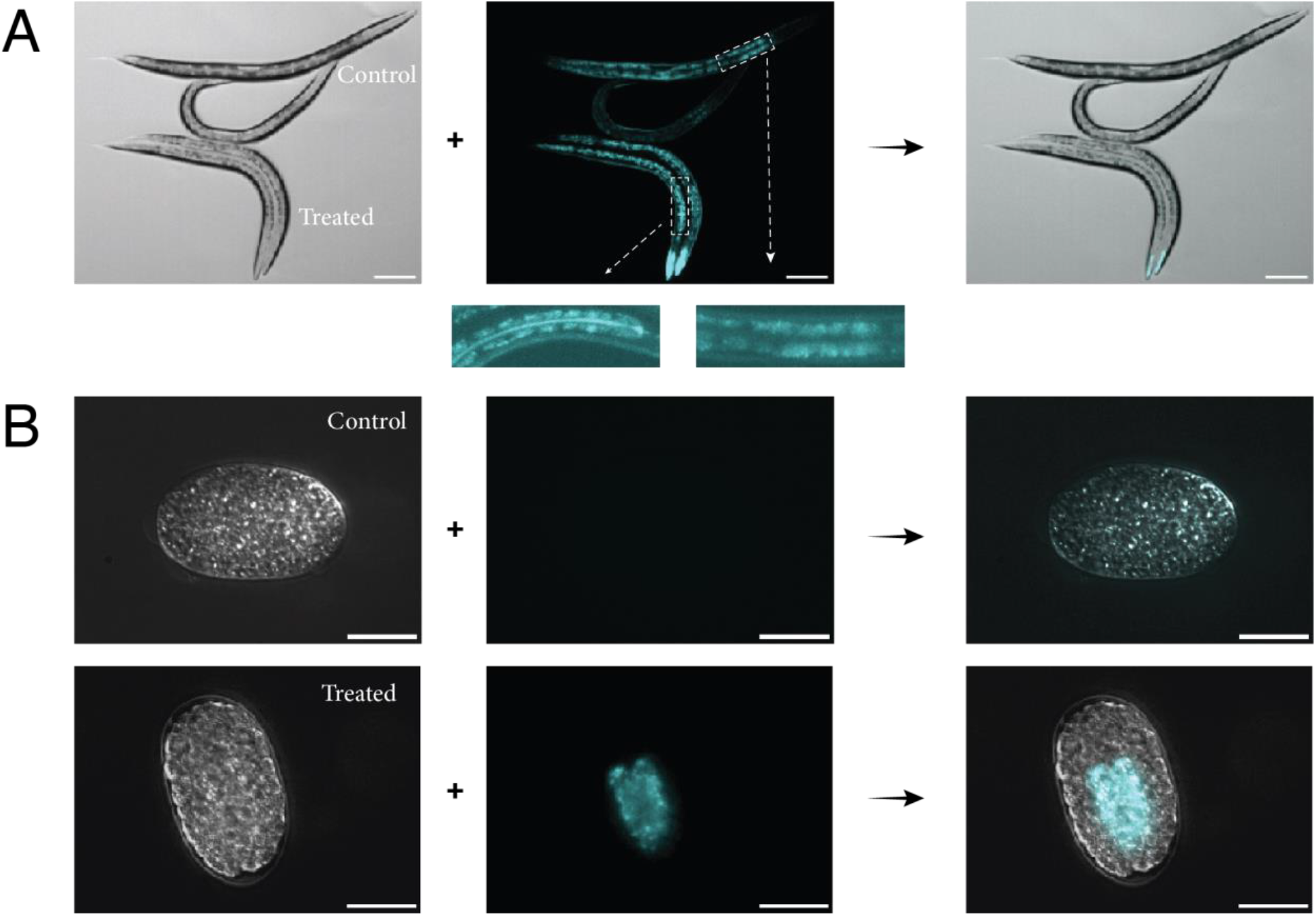
Distribution of fluorescently-labeled avocadyne in C. elegans larvae and embryos. (A) Representative images of control and treated L4 larvae: DIC (left), fluorescence (center), and merged DIC/fluorescence (right). In any panel of the row, the control and treated samples are paired and labeled. (B) Representative images of control and treated embryos: DIC (left), fluorescence (center), and merged DIC/fluorescence (right). Control samples were always obtained by treating worms or embryos with DMSO solution of 3-azido-7-hydroxycoumarin that has no fluorescence. Treated samples were always obtained by treatment with hydroxycoumarin-tagged avocadyne whose fluorescence was detected at 480 nm following excitation. The weak fluorescence observed for the control in the central panel of row A arises from L4 autofluorescence background. Scale bars: 100 µm (A); 20 µm (B).

In addition to the distribution described above, treated L4 larvae exhibited avocadyne-conjugate fluorescence in a population of normal-sized LD distinct from the DHS-3::GFP-positive LD that mark conventional neutral lipid stores (Fig. 6D). Treatment with fluorescently labeled avocadyne increased the total number of LD by 73% relative to controls. Occasional colocalization of the avocadyne conjugate with DHS-3::GFP was also observed, together with a subpopulation of enlarged LD labeled only by DHS-3::GFP (Fig. S7). The exclusion of the avocadyne conjugate from these enlarged LD suggests that they may arise through the fusion of pre-existing, unlabeled LD. Thus, the composite LD profile shown in Figure S7 and the simpler one devoid of enlarged LD of Figure 6D may be qualitatively consistent with differences in metabolic response enforcing lipid storage both by LD fusion and LD proliferation, rather than the latter only [38–40, 46–49].

**Figure 6.**
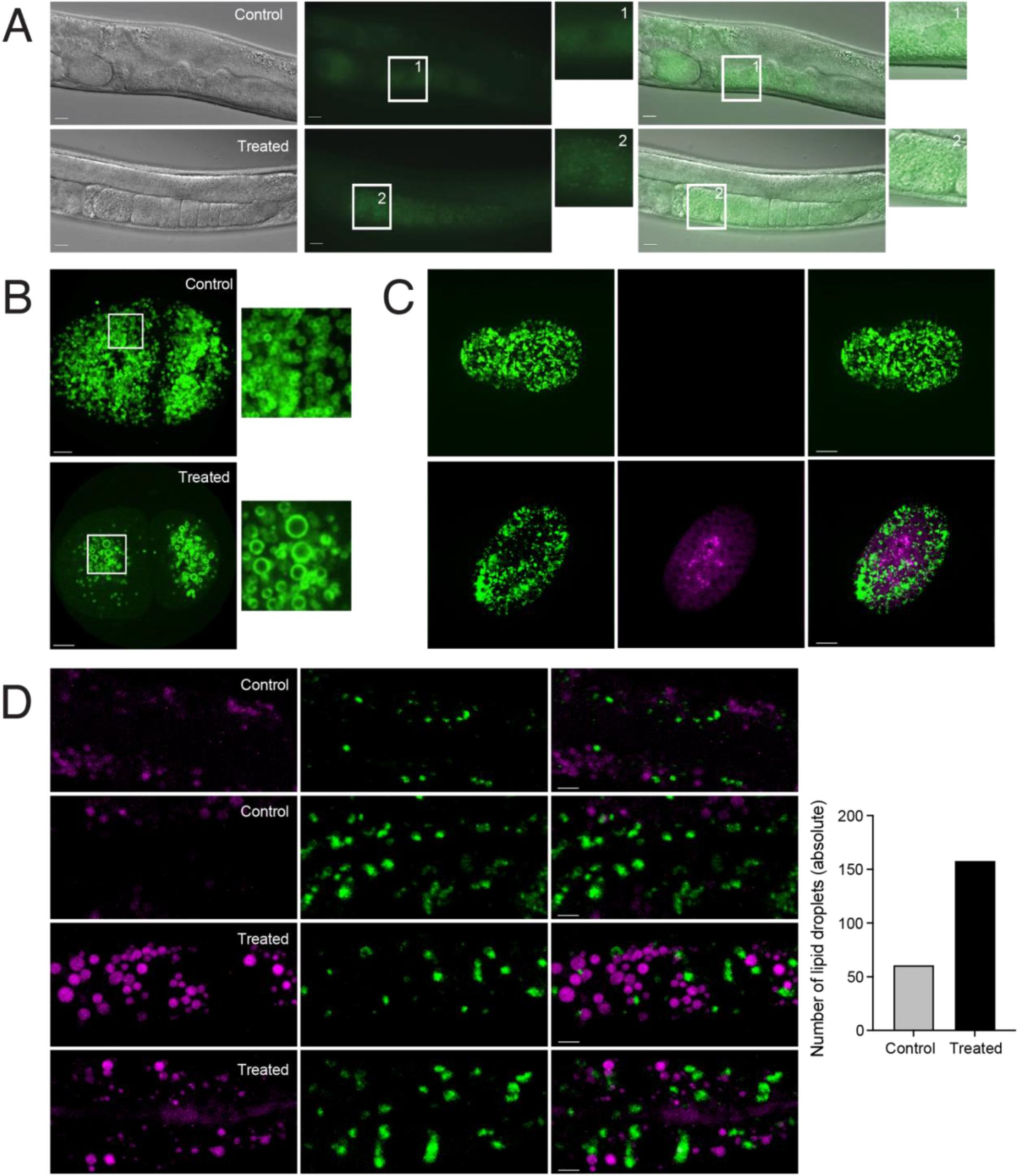
Lipid BODIPY™ staining, GFP-labeled LD and coumarin-conjugated avocadyne fluorescence imaging of C. elegans larvae and embryos. (A) Images of control (DMSO) and avocadene-acetate-treated L4 larvae: DIC (left), BODIPY™ staining (center), DIC/BODIPY™ merging (right). The magnified insets highlight the lipid droplets (LD) difference between treated and control L4 samples. (B) GFP-labeled LD confocal microscopy images of control (DMSO) and avocadene-acetate-treated embryos, strain VS50 plin-1(hj178[PLIN-1::GFP]). (C) Confocal microscopy images of GFP-tagged LD and avocadyne conjugate fluorescence from control (DMSO) and treated (hydroxycoumarin-conjugated avocadyne in DMSO) embryos, strain VS50 plin-1(hj178[PLIN-1::GFP]). The magenta pseudo-color (center and right) highlights the tagged avocadyne fluorescence, present only in the treated sample (excitation at 405 nm, emission in the interval 480-502 nm). The green color (left and right) represents the GFP fluorescence (excitation 488 nm, emission 507 nm). Merging of magenta and green fluorescence is shown in the right column. (D) Representative cross-section combinations from confocal microscopy images of control (DMSO) and treated (hydroxycoumarin-conjugated avocadyne in DMSO) GFP-labeled-LD [LIU1 (dhs-3p::DHS-3::GFP)] L4 larvae intestinal region. The magenta pseudo-color (left and right) highlights the tagged avocadyne fluorescence, present only in the treated sample, and the gut granules autofluorescence, present also in the control sample (excitation at 405 nm, emission in the interval 480-502 nm). The green color (center) represents the GFP fluorescence (excitation 488 nm, emission 507 nm). Merging of magenta and green fluorescence is shown in the right column. The lateral bar plot reports the LD counting across all the control and treated sample images. Scale bars: 50 µm (A); 10 µm (B); 10 µm (C); 10 µm (D).

Affinity-driven interactions between coumarin-tagged avocadyne and TG chains (SFA, MUFA and flexible PUFA) could facilitate incorporation of the conjugate into nascent LD, accounting for the detectable colocalization in some normal-sized droplets (Fig. S7). In the enlarged DHS-3::GFP-positive LD, however, the abundance of neutral lipids may dilute the conjugate signal below the detection threshold. Although LD fusion and proliferation are regulated by distinct biochemical signals [49], the two responses may overlap under our experimental conditions. Consistent with this interpretation, weak colocalization was also detectable in the proliferation-dominant profile shown in Figure 6D (see Fig. S8).

Some autofluorescent circular structures were visible in the DMSO controls (Fig. 6C) and DMSO-plus-hydroxycoumarin controls (not shown), particularly in intestinal gut granules, which are known to exhibit intrinsic fluorescence across a broad range of wavelengths. In contrast, avocadyne-conjugate fluorescence appeared ring-like, outlining hollow vesicular structures rather than the solid puncta observed in controls (Fig. S9). BODIPY™ staining of control and avocadene-acetate-treated L4 larvae likewise revealed a shift from a diffuse distribution in control animals to a more concentrated, punctate pattern following treatment (Fig. 6A), consistent with an increase in LD number.

Following treatment with the fluorescent avocadyne derivative, *C. elegans* embryos exhibited a distinct fluorescence signal upon excitation at 350 nm, demonstrating internalization of the conjugate compound. Differential interference contrast (DIC) and fluorescence microscopy revealed intracellular accumulation of the conjugate in embryos, accompanied by developmental arrest and eventual lethality (Fig. 5B). Because the eggshell normally forms a strong permeability barrier to many small molecules, the ability of avocadyne and its hydroxycoumarin conjugate to cross the eggshell is noteworthy, although the predicted reduction in conjugate solubility may limit its permeability (Fig. S3). Mitochondrial disruption likely contributes substantially to the observed lethality, given the critical role of mitochondrial function during early *C. elegans* embryogenesis. By contrast, inhibition of ACC/POD-2 is generally associated with defects in embryonic polarity and osmotic sensitivity [50–52], bearing therefore a different role and relevance with respect to the larval states.

NMR-based lipidomic analysis revealed modest increases in lipid abundance in embryos treated with avocadene acetate (Fig. 1E). Compared with the corresponding profile in L4 larvae, however, the markedly smaller increase in PC and the still smaller increase in PL (the latter comparable to the average increase among remaining lipid species) are consistent with lipid storage through enlargement or fusion of existing LD rather than the formation of new LD [38–40, 46–49]. Accordingly, embryos treated with hydroxycoumarin-tagged avocadyne (Fig. 5B), as well as embryos expressing GFP-tagged perilipin and treated with either avocadene acetate (Fig. 6B) or conjugated avocadyne (Fig. 6C), displayed substantially larger LD than those observed in L4 larvae. As in L4 larvae, limited colocalization of neutral lipids with conjugated avocadyne was also observed (Fig. S8), which became more pronounced at later embryonic stages (not shown). Taken together, the enlarged droplets and pronounced reduction in LD number support a shift toward LD fusion and increased lipid storage in treated embryos. These findings suggest that avocadene/yne compounds impair normal embryogenesis through biochemical mechanisms distinct from those predominating in L4 larvae, without producing obvious defects in polarity or osmotic sensitivity associated with *pod-2* loss of function [50–52].

## Discussion

### Avocado-derived compounds induce LD profile alteration and accumulation of lipids in C*. elegans*

In this study we investigated the effects at the lipid level of avocado-derived bioactive compounds avocadene, avocadyne, and their acetate derivatives on lipid composition and abundance in *C. elegans* embryos and L4 larvae, by combining NMR spectroscopy and lipid profiling with click-chemistry-based fluorescent labeling and microscopy. Building upon our previous work [7–9], we sought not only to confirm the internalization and metabolic incorporation of those compounds, but also to examine their selectivity and potential interactions with specific lipid classes and characterize their broader effects on lipid profiles. Treatment produced an appreciable and consistent increase in lipid content in both embryos and L4 larvae, indicating altered lipid storage and disruption of lipid homeostasis at both development stages. In L4 larvae, the increases in phospholipids (PL) and phosphatidylcholines (PC) were particularly pronounced, whereas triglycerides (TG) and other lipid classes increased more modestly. By contrast, embryos showed substantially smaller increases in PC and PL, with the increase in PL comparable in magnitude to those observed for the other lipid classes, including TG.

The observed lipid accumulation also connects with the avocadene/yne derivative interactions inferred from diffusional properties. NMR DOSY analysis revealed markedly lower diffusion coefficients for avocadene acetate in extracts from treated embryos and larvae than for the isolated compound. Initially interpreted primarily as evidence that avocadene acetate retained its molecular integrity following internalization and extraction, the measurements reported in Table 1 acquire additional significance when considered alongside the lipidomic data. In the chloroform fraction of the Folch extracts, avocadene acetate appears not to diffuse freely but instead to associate dynamically with larger lipid species or assemblies, likely through their shared hydrophobic character. Given the composition of these extracts, the reduced diffusivity is consistent with interactions involving neutral-lipid or membrane-associated lipid pools. Although these *ex vivo* measurements do not directly demonstrate association in intact animals, they support the possibility that similar interactions occur in vivo.

To test this inference, we first established a click-chemistry labeling strategy by reacting an equimolar mixture of avocadyne and avocadyne acetate with 3-azido-7-hydroxycoumarin through copper-catalyzed azide–alkyne cycloaddition (CuAAC) (Fig. 2A). Formation of the fluorescent triazole conjugates was confirmed by UV fluorescence and NMR, demonstrating that both the non-acetylated and acetylated forms could be labeled for subsequent visualization and tracking. We then treated C. elegans embryos and L4 larvae with the clickable avocadyne compounds, extracted the lipids, and fluorescently labeled the incorporated compounds by CuAAC. This approach allowed us to trace the avocadyne derivatives and isolate lipid species associated with them.

Fluorescence-guided selective TLC separation followed by NMR analysis revealed the association with SFA, MUFA and PUFA chains containing two or more methylene groups separating successive double bonds. These findings support the preferential association of non-acetylated and acetylated avocadyne with SFA, MUFA, and conformationally flexible PUFA chains, predominantly esterified to glycerol within TG, and suggest similar behavior for the corresponding avocadene derivatives. This selectivity may reflect differences in acyl-chain conformational dynamics: closely spaced double bonds restrict conformational sampling and may prevent optimal packing against the flexible avocadyne or avocadene chain. PUFA with more widely spaced double bonds, however, may retain sufficient flexibility to maximize hydrophobic contacts while incurring a smaller conformational-entropy penalty upon association. This balance could potentially favor their association even over that of SFA and MUFA.

The detection of TG signatures in the TLC-resolved fluorescent fractions is particularly noteworthy. Because TG are the major form of lipid storage, their association with fluorescent avocadyne conjugates suggests a favorable interaction between these compounds and neutral lipid pools. Consistent with this interpretation, lipidomic analysis following avocadene acetate exposure revealed an appreciable increase in total lipid content, including TG. The storage response differed markedly between developmental stages: embryos contained fewer, enlarged LD, consistent with droplet fusion, whereas L4 larvae exhibited an increase in LD number, consistent with droplet proliferation. In larvae, the disproportionately larger increases in PC and PL relative to TG further support the formation of new LD, as an increased number of small droplets requires a greater monolayer surface area relative to the volume of the neutral-lipid core [38–40, 46–49]. These newly formed LD may incorporate avocadene or avocadyne derivatives and associated neutral lipids, whereas changes in conventional, pre-existing neutral-lipid LD may represent a downstream metabolic response rather than direct compound accumulation.

The enrichment of MUFA and conformationally flexible PUFA in larval extracts, together with their association with fluorescently labeled avocadyne, further suggests that these compounds influence the composition and metabolism of energy-dense lipid stores. If incorporated into LD monolayer phospholipids, these unsaturated acyl chains could increase monolayer fluidity [45] and facilitate the high curvature required to form numerous small LD. In embryos, analogous changes in membrane fluidity could conceivably contribute to the osmotic-sensitivity phenotypes associated with inhibition of ACC/POD-2 [7–9, 50–52]. Besides the limited embryonic increase of UFA levels, however, the present study provides no evidence for this possibility, in line with the absence of any polarity defects in embryonic development after avocadene/yne species treatment.

Overall, the lipid flux remodeling observed in embryos and L4 larvae, likely triggered by inhibition of ACC/POD-2 by avocadene/yne compounds, may represent an adaptive response to lipotoxic shock under conditions of impaired mitochondrial function [38, 40, 53, 54]. In embryos, the shift toward increased lipid storage appears insufficient to restore lipid homeostasis and support continued development [52]. In L4 larvae, by contrast, lipid trafficking is redirected toward the production of numerous LD following transient lipotoxicity-induced paralysis. This response may help sequester excess lipids and mitigate lipotoxic damage.

LD are now widely recognized as active organelles that buffer metabolic and oxidative stress [38–41, 55]. In many systems, stress induces LD formation sequesters potentially harmful lipid species to limit lipotoxicity, protect against ROS and ER (endoplasmic reticulum) stress, support mitochondrial function, and coordinate lipid traffic through contact sites with mitochondria and other organelles [38, 39, 41, 53, 54, 56, 57].

### Avocadyne localization in larval and embryonic tissues

The fluorescent avocadyne conjugate was used as a proxy to examine the distribution of the avocadene- and avocadyne-derived compounds in *C. elegans* embryos and L4 larvae. Fluorescence microscopy and spectroscopy indicated that the conjugate was readily taken up and internalized by C. elegans, exhibiting distinct tissue- and stage-specific localization patterns. In L4 larvae, the rapid appearance of fluorescence in the pharynx and its subsequent accumulation in the intestinal lumen were consistent with ingestion through feeding and passage through the digestive tract. Retention in the pharyngeal isthmus and downstream intestinal regions may reflect transient association with the musculature, secretory structures, or other components of the digestive system. This distribution is particularly relevant given the central role and high metabolic rate of the pharynx during nutrient intake and suggests a potential route through which avocadyne could affect digestive physiology.

In embryos, fluorescently tagged avocadyne crossed the eggshell, which typically forms a robust permeability barrier, and accumulated within embryonic cells. Its penetration nevertheless appeared less efficient than that of unconjugated avocadyne, consistent with the increased polar surface and reduced solubility predicted for the conjugate (Fig. S3). Given the high energy demand of rapidly dividing embryonic cells [50–52], the disruption of mitochondrial integrity by avocadene/yne derivatives likely underpins the observed development arrest and lethality.

## Conclusions

Biochemical evidence confirming *C. elegans* ACC/POD-2 as a primary target [9] and partial rescue by malonyl-CoA [8] functionally link the biological effect of avocadene/yne free alcohols and acetates to reduced malonyl-CoA production following ACC/POD-2 inhibition. ACC/POD-2 catalyzes the rate-limiting step for malonyl-CoA production, which both supports *de novo* fatty-acid synthesis and restricts mitochondrial β-oxidation through inhibition of CPT-I [30–34]. ACC inhibition should therefore disrupt the balance between fatty-acid biosynthesis and mitochondrial oxidation, increasing the burden of fatty acids and their catabolites on mitochondria and thereby inducing metabolic stress [33, 34, 50–52, 58, 59].

Consistent with this model, treatment with avocadene/yne-derived compounds induced severe mitochondrial dysfunction, elevated ROS, and transient paralysis in *C. elegans* L4 larvae [8]. In embryos, where ACC/POD-2 also regulates development polarity and osmotic sensitivity [50–52], mitochondrial dysfunction following treatment instead led to developmental arrest [8].

Despite ACC inhibition, lipidomic and NMR analyses of treated L4 larvae revealed an increase in total lipid content, including a pronounced rise in PL levels, particularly PC, and more modest increases in MUFA and PUFA chains, predominantly esterified as TG. Similar paradoxical lipid increments have previously been reported [35–37] and attributed to secondary effects involving activation of anaplerotic pathways. Because LD biogenesis increases the monolayer surface area relative to the volume of stored neutral lipids, it requires increased phospholipid production, particularly PC [47–49]. LD are the principal organelles for neutral-lipid storage and are surrounded by a phospholipid monolayer in which PC is the predominant lipid [38–41, 47–49]. Thus, the pronounced PC increase relative to TG accumulation in avocadene-acetate-treated larvae is consistent with extensive LD biogenesis and expansion of the total LD surface area. In contrast, the substantially smaller increase in PC in treated embryos, which was comparable to the increase in TG, favors expansion of existing LD volume through droplet fusion rather than the formation of numerous new LD in response to a higher lipid storage demand [38–41, 47–49]. Together, these findings suggest that ACC/POD-2 inhibition and mitochondrial impairment drive an adaptive LD response in avocadene/yne-treated *C. elegans* L4 larvae, in which a new LD subpopulation is formed in the attempt to detoxify the organism from both excess fatty acids and the drug itself [38, 39, 53, 54]. Sequestration of avocadene/yne-derived compounds within these droplets could lower the effective free concentration of the compound at mitochondria [33, 34], potentially explaining the transient nature of paralysis at sublethal doses even though lipid composition and LD profile remain altered. In embryos, the same perturbation enhances neutral-lipid storage and LD expansion but fails to restore viability, possibly because embryos have less capacity to deploy compensatory metabolic pathways. As a result, treated embryos undergo developmental arrest and fail to hatch despite retaining normal early cell polarity.

## Abbreviations

ACC: Acetyl-CoA Carboxylase
*C. elegans*: *Caenorhabditis elegans*
CPT-I: Carnitine Palmitoyl Transferase I
DG: *1,2* or *1,3* diacyl-glycerides
DHS-3: short chain Dehydrogenase/reductase
DHSA: Dihydrosterculic Acid
DIC: Differential Interference Contrast
DOSY: Diffusion Ordered Spectroscopy
*E. coli* OP50: *Escherichia coli* strain OP50
FA: Fatty Acids
FID: Free Induction Decay
GFP: Green Fluorescent Protein
HSQC: Heteronuclear Single Quantum Coherence
INEPT: Insensitive Nuclei Enhanced by Polarization Transfer
LD: Lipid Droplets
MUFA: Monounsaturated Fatty Acids
NMR: Nuclear Magnetic Resonance
NOESY: Nuclear Overhauser Effect Spectroscopy
PC: Phosphatidylcholines
PE: Phosphatidylethanolamines
PL: Phospholipids
PLIN: Perilipin
PUFA: Polyunsaturated Fatty Acids
ROS: Reactive Oxygen Species
SFA: Saturated Fatty Acid
TG: Triglycerides
TLC: Thin Layer Chromatography
TOCSY: Total Correlation Spectroscopy
UFA: Unsaturated Fatty Acids
Δ: Unsaturation
*ω3, ω6, ω9*: Unsaturation position of fatty acids with respect to terminal (ω) methyl

## Supplementary Information

The online version contains supplementary material available on line.

## Acknowledgements

The institutional supports of the NYUAD Core Technology Platform, for instrumentation, and Mrs. Perihan Bermamet, for administration, are gratefully acknowledged. We also thank Ashwini Dolle for the help with some panels.

## Author contributions

Investigation: YH, GE, SG, FSR, RH, YM & LA. Methodology: YH, GE & HZF. Formal analysis: YH, GE, SG, FSR, RH, YM & LA. Resources: KCG & FP. Validation: YH, GE, SG, FSR, RH & HZF. Supervision: GE, HZF, KCG & FP. Conceptualization: GE, YH, HZF, KCG & FP. Project administration: GE & HZF. Funding acquisition: KCG & FP. Writing-original draft. YH & GE. Writing-review and editing. YH, GE, SG, FSR, RH, KCG & FP.

## Availability of data and materials

The datasets supporting the conclusions of this article are included within the additional file of Supporting Information. The remaining datasets used and/or analyzed during the current study are available from the corresponding author on reasonable request.

