## Supplementary Information for "Avocado-derived compounds alter lipid homeostasis and lipid droplets profile in *Caenorhabditis elegans*"

[1] Science Division, Biology Program, New York University Abu Dhabi, Abu Dhabi, United Arab Emirates. [2] Center for Genomics and Systems Biology, New York University Abu Dhabi, Abu Dhabi, United Arab Emirates. [3] Dipartimento di Medicina, Università di Udine, 33100 Udine, Italy. [4] Core Technology Platform, New York University Abu Dhabi, Abu Dhabi, United Arab Emirates. [5] Center for Genomics and Systems Biology, Department of Biology, New York University, New York, NY, USA. [6] Istituto Nazionale Biostrutture e Biosistemi, 00136 Rome, Italy.

*Correspondence
Gennaro Esposito
Fabio Piano

Hala Zahreddine Fahs

Kristin C. Gunsalus


### SUPPLEMENTARY INFORMATION

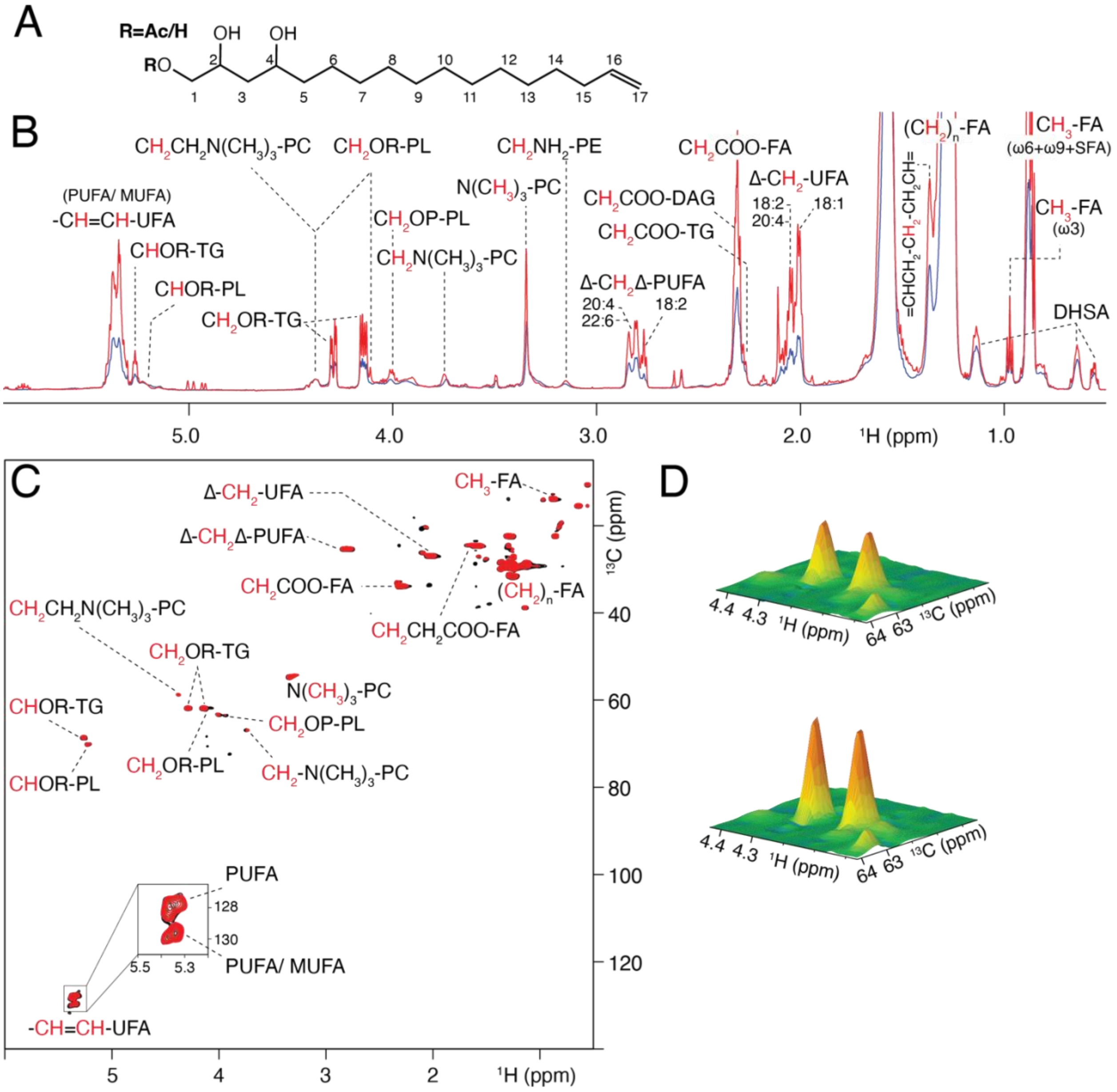


**Figure S1**. NMR analysis comparison of lipid profiles in control and avocadene acetate-treated *C. elegans* embryos. (A) Chemical structure of avocadene (R=H) and avocadene acetate (R=Ac). (B) Overlay of 1D $^{1}$H NMR spectra from control (blue) and treated (red) embryos samples with lipid peak assignments. Contributing protons are marked in red. Abbreviations: FA, fatty acids; UFA, unsaturated fatty acids; MUFA, monounsaturated fatty acids; PUFA, polyunsaturated fatty acids; TG, triglycerides; PL, glycerophospholipids; PC, phosphatidylcholines; PE, phosphatidylethanolamines; DAG, 1,2 or 1,3 diacyl-glycerides; DHSA, dihydrosterculic acid; SFA, saturated fatty acid; Δ, unsaturation; w3, w6, w9 = unsaturation position with respect to terminal methyl. (C) 2D $^{13}$C–$^{1}$H HSQC spectra of lipid extracts of embryos; control (red) and treated (black). Peak assignments are indicated, with the contributing proton(s) and carbon(s) highlighted in red.

(D) 3D representation of the $^{13}C$–$^{1}H$ HSQC cross-peaks of the triglyceride glycerol methylenes ($CH_2OR$-TG) showing the volume difference between control (top) and treated (bottom) embryo samples.

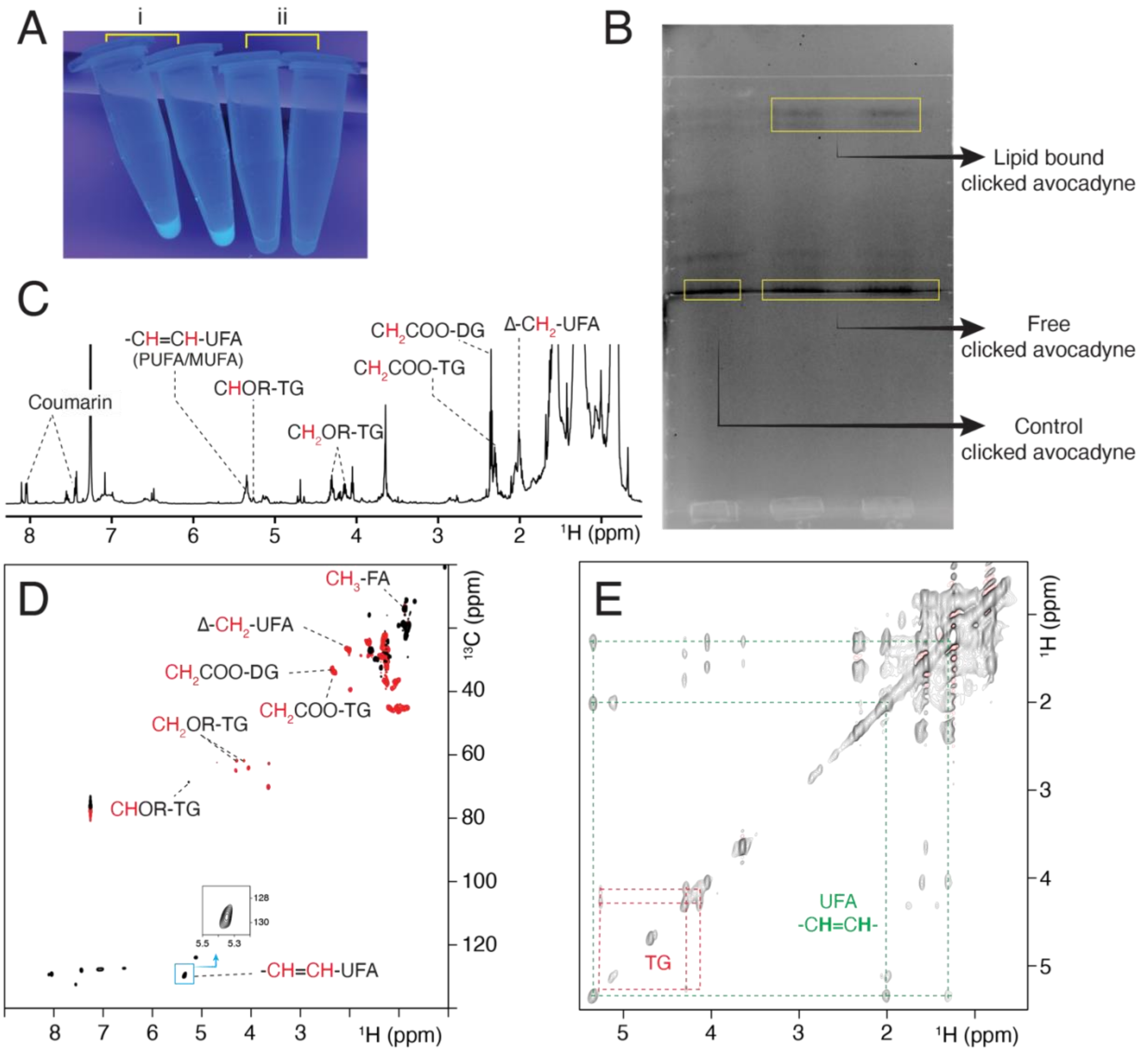


**Figure S2**. Fluorescence and NMR tracing of avocadyne-treated embryo extract after fluorophore conjugation and TLC separation. (A) UV lamp irradiation (365 nm) of avocadyne-treated C. elegans L4 larvae suspensions before (i) and after (ii) conjugation to the coumarin derivative fluorophore (B) TLC development highlighting fluorescent strip separation corresponding to free and lipid-bound hydroxycoumarin-tagged avocadyne. (C)$^1$H 1D spectrum in $CDCl_3$ of the Folch extract from avocadyne-treated larvae after submission to coumarin derivative conjugation, TLC resolution and chloroform extraction selectively carried out on fluorescent TLC bands. The lipid assignments are reported. (D) $^{13}$C-$^1$H edited HSQC spectrum showing the assignments of the relevant lipid signals. The insets indicate the resonances corresponding to the UFA olefinic nuclei (CH=CH-UFA) that were observed in different samples, namely lipids extracted from fluorescent TLC bands (black), lipids from untreated embryos (blue), and lipids from avocadyne-treated embryos (purple). (E) $^1$H-$^1$H TOCSY spectrum highlighting triglycerides (TG) and unsaturated fatty acids (UFA) connectivities.

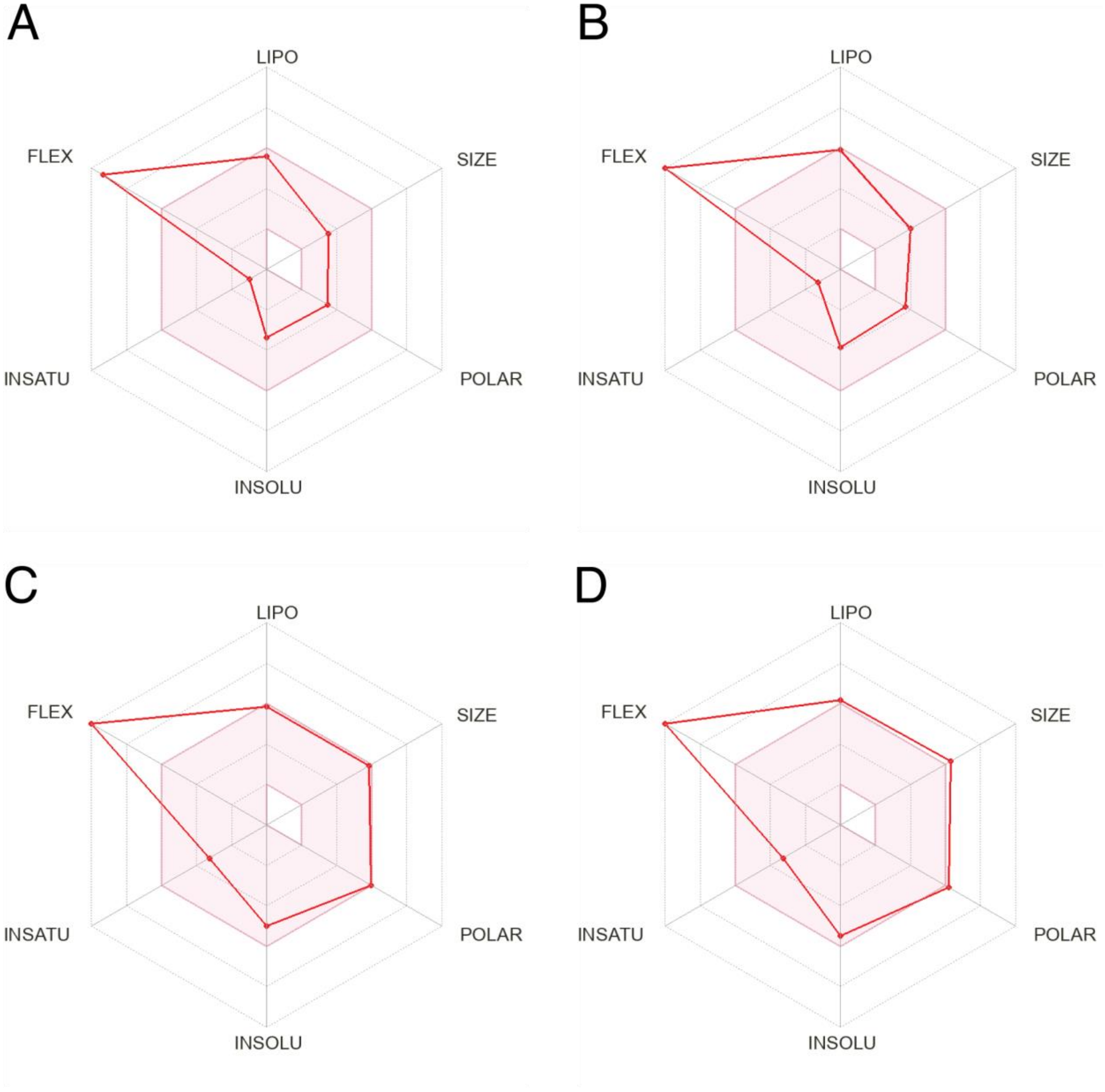


**Figure S3**. Chemical characterization and comparative analysis of avocadyne, avocadyne acetate, and their coumarin conjugates. SwissADME physicochemical characterization of (A) avocadyne, (B) avocayne acetate, (C) avocadyne-coumarine conjugate, (D) avocadene acetate-coumarine conjugate. LIPO: lipophility, SIZE: size of the molecule, PLOAR: polarity, INSOLU: insolubility, INSATU: insaturation, FLEXL: flexibility.

The chemical characterization of avocadyne, avocadyne acetate, and their coumarin-linked conjugates was performed using SwissADME [45] to evaluate and compare their physicochemical properties. Avocadyne exhibited a topological polar surface area (TPSA) of 60.69, a molar refractivity of 85.48, and a log P value of 3.09, indicating moderate lipophilicity, and balanced hydrophilic-hydrophobic properties. Avocadyne acetate showed a slightly higher TPSA of 66.76, a molar refractivity of 95.22, and a log P of 3.67, suggesting enhanced polarizability and lipophilicity compared to avocadyne. These differences may influence their respective solubility, permeability, and pharmacokinetics. The coumarin-linked conjugates exhibited distinct physicochemical characteristics due to the addition of the coumarin moiety. The avocadyne-coumarin conjugate had a TPSA of 128.70, molar refractivity of 137.32, and a log P of 3.55, reflecting a significant increase in polar surface area and molecular size while maintaining moderate lipophilicity. Similarly, the avocadyne acetate-coumarin conjugate showed a TPSA of 134.77, molar refractivity of 147.06, and a log P of 4.12, indicating the highest lipophilicity and molecular size among all considered species. Comparatively, the coumarin-linked conjugates have much larger polar surface areas and molecular sizes than their parent compounds accompanied by increased insolubility, which may enhance their interaction potential with polar environments but could reduce permeability.

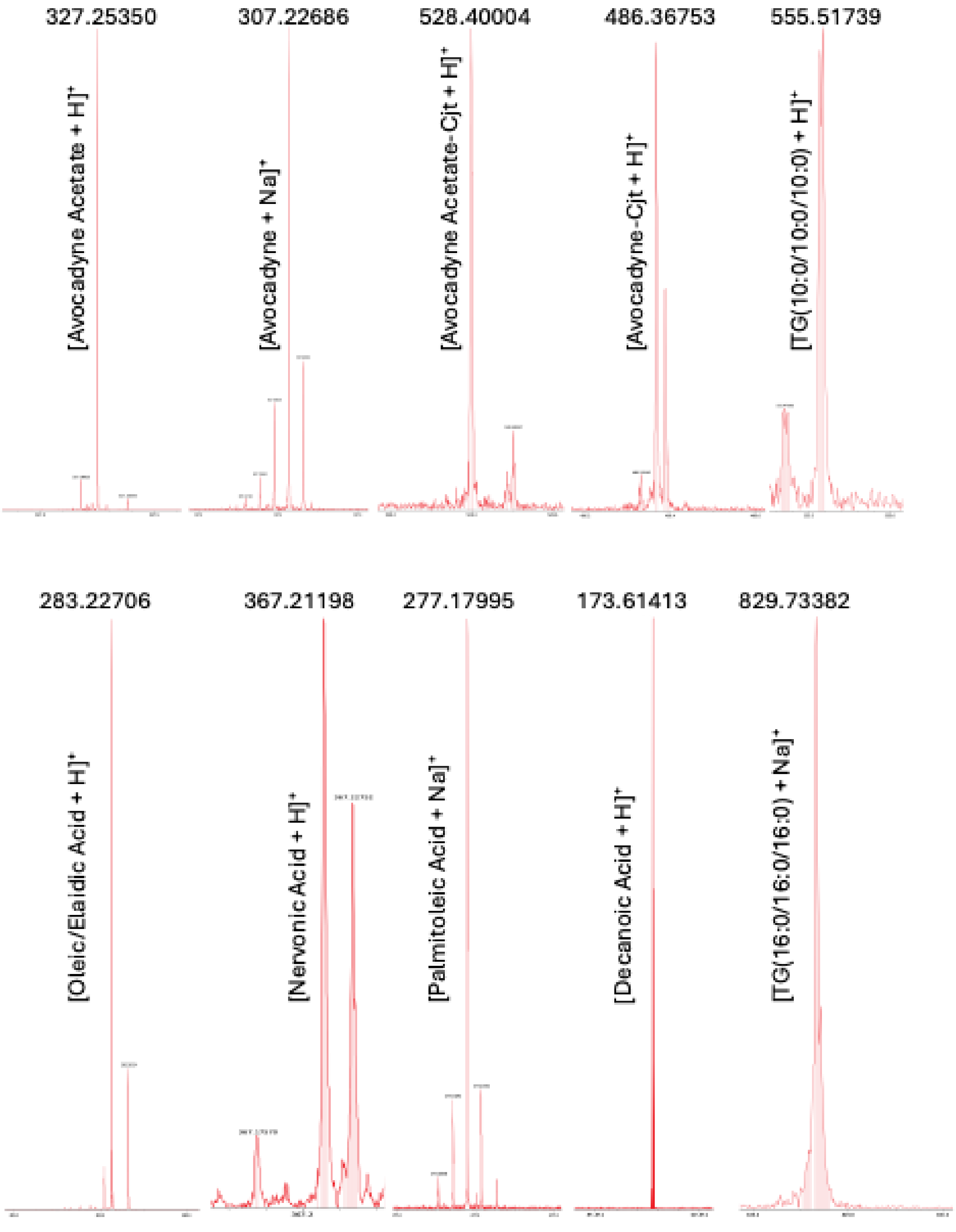


**Figure S4**. Mass spectrometry analysis of TLC selective extraction from C. elegans embryos. The most prominent signals (single-charge cations) are reported with the mass-over-charge value (in Th units) of the corresponding proton or sodium adduct.

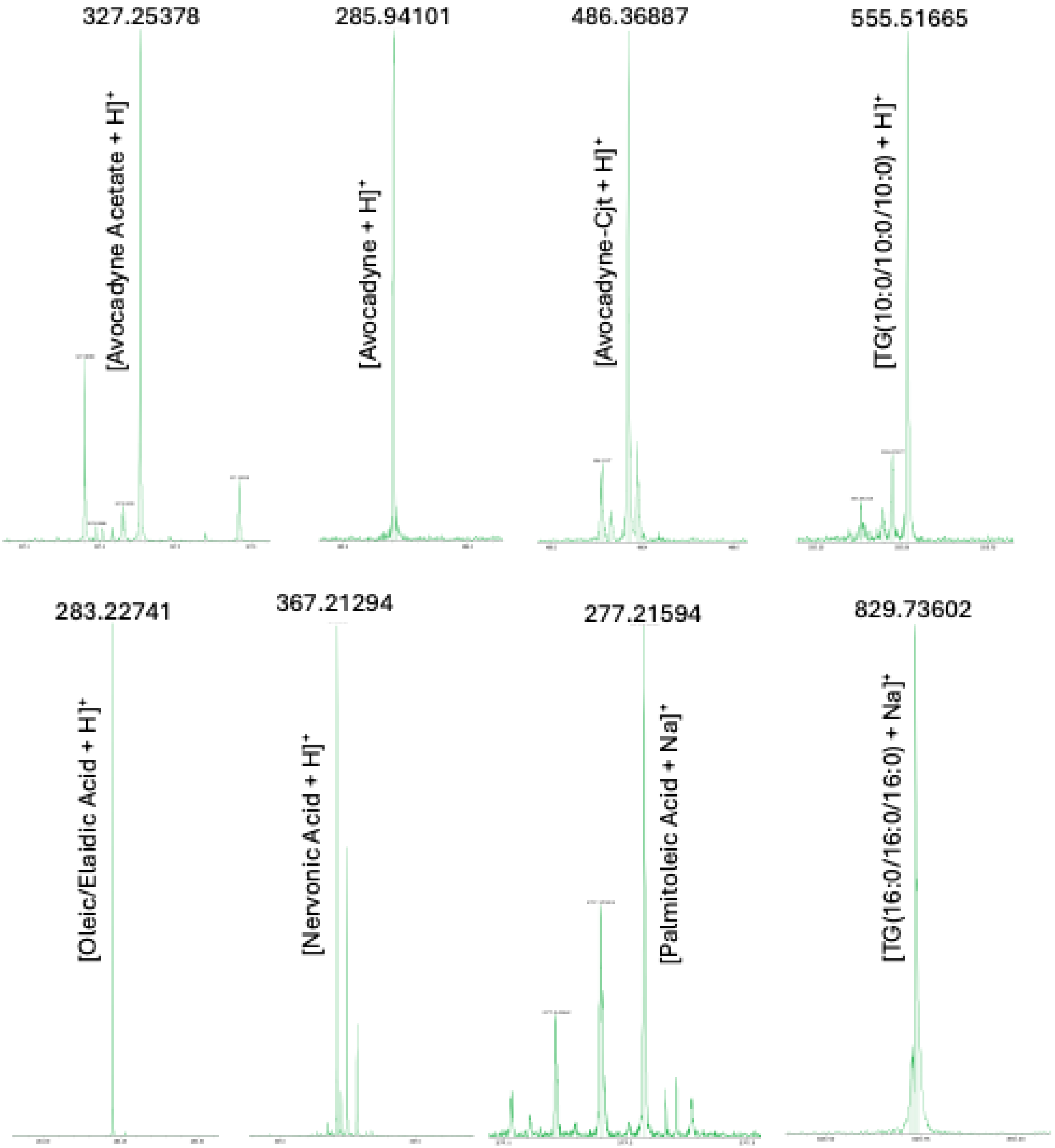


**Figure S5**. Mass spectrometry analysis of TLC selective extraction from C. elegans L4 larvae. The most prominent signals (single-charge cations) are reported with the mass-over-charge value (in Th units) of the corresponding proton or sodium adduct.

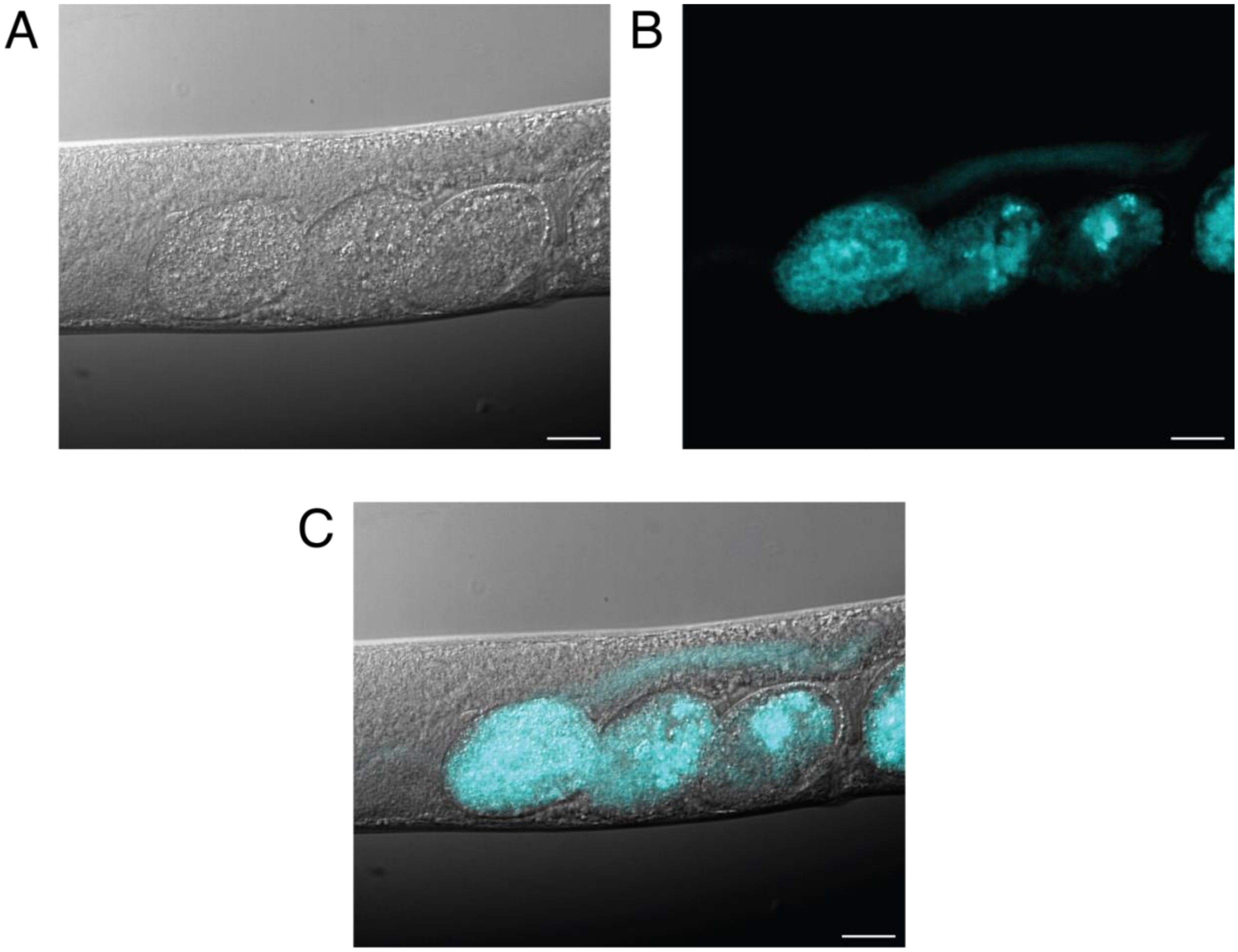


**Figure S6**. Distribution of fluorescently-labeled avocadyne derivatives in C. elegans larvae. Representative images of treated gravid adults showing the diffusion of tagged avocadyne derivatives into embryos (A) DIC, (B) Fluorescent image, (C) overlay of A and B. Scale bars: 20 µm

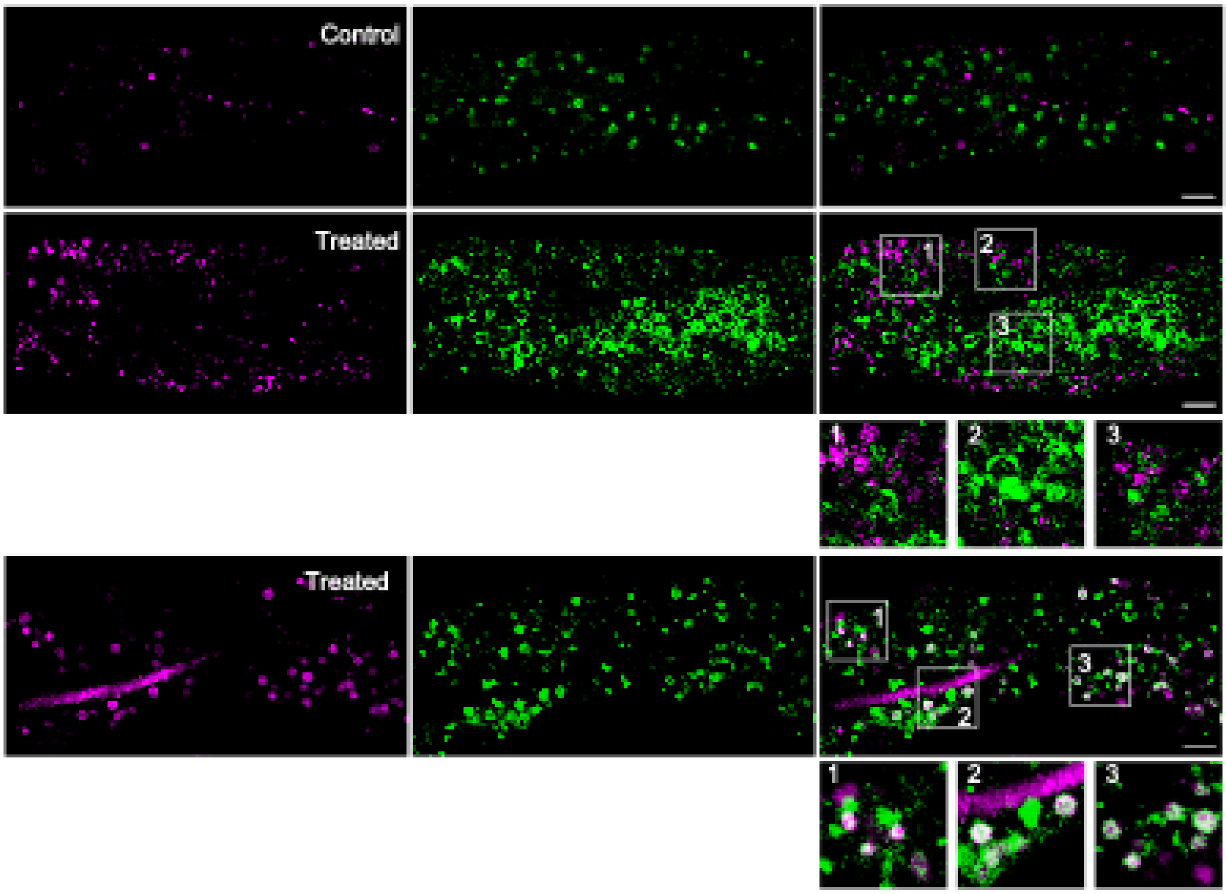


**Figure S7**. Representative confocal images of control (DMSO) and treated (hydroxycoumarin-conjugated avocadyne in DMSO) GFP-labeled LD [LIU1 (dhs-3p::DHS-3::GFP)] L4 larvae intestinal region. Subpopulations of enlarged-volume LD (treated upper panels) and regular-size LD (treated lower panels) are displayed, along with the separation or colocalization of their different fluorescence. The magenta-highlighted continuous segment in the regular LD panels arises from the presence of conjugated avocadyne in the intestinal lumen. The magenta pseudo-color (left and right) highlights the tagged avocadyne fluorescence (present only in the treated samples) and the gut granules autofluorescence (present also in the controls sample) (excitation at 405 nm, emission in the interval 480-502 nm). The green color (center and left) represents the GFP fluorescence (excitation 488 nm, emission 507 nm). Merging of magenta and green fluorescence is shown in the right column, where pale grey identifies colocalization. Scale bars: 10 µm.

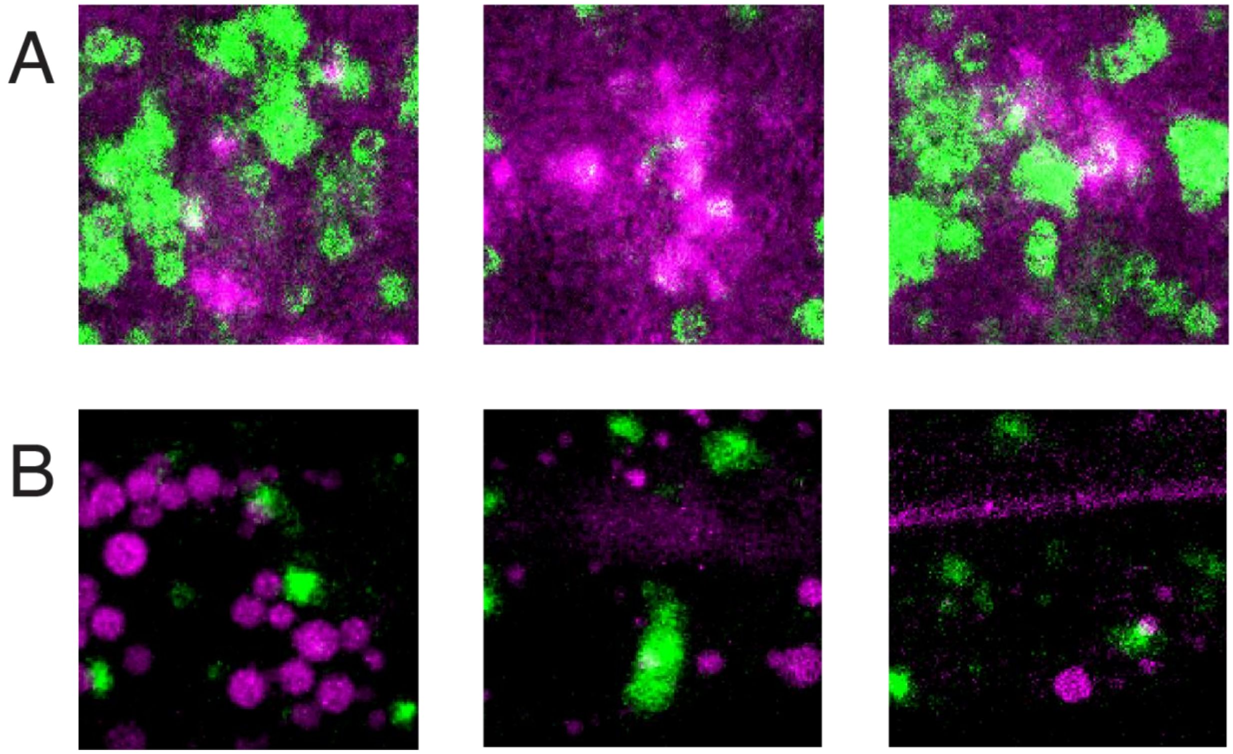


**Figure S8**. Details from f the same confocal microscopy image as reported in Figure 6C (A), and Figure 6D (B) of main text. Colocalization of avocadyne-conjugate and GFP-labeled neutral-lipid LD fluorescence can be inferred from the light grey regions.

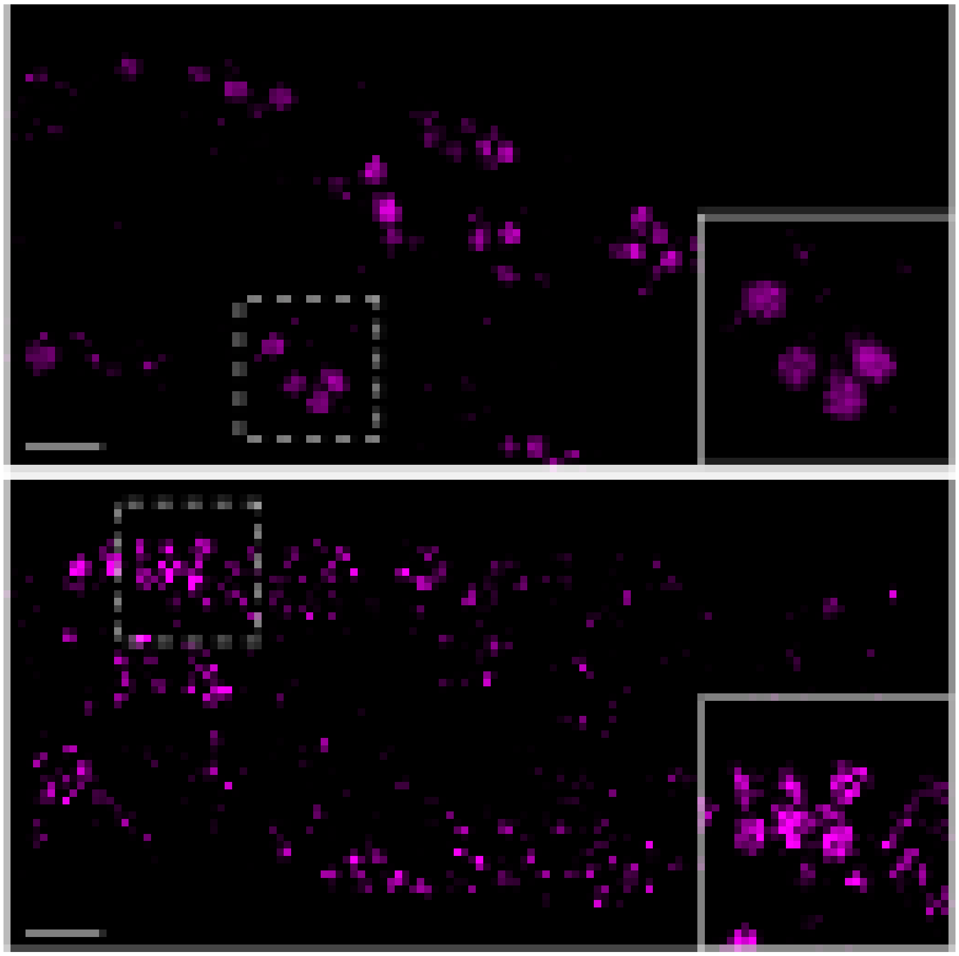

**Figure S9**. Hydroxycoumarin-conjugated avocadyne derivative fluorescence imaging in C. elegans larvae. Representative confocal images of control (DMSO, upper panel), and treated (hydroxycoumarin-conjugated avocadyne in DMSO, lower panel) L4 larvae intestinal region. The magenta pseudo-color highlights the gut granules autofluorescence and the LD tagged avocadyne fluorescence. The fluorescence signals from the avocadyne conjugates appear as hollow vesicles rather than solid spheres as observed for the gut granules in control images. Scale bars: 10 μm for both upper and lower panels.

**Table S1**: List of lipids identified based on COLMAR NMR online lipid server predictions and manual analysis by referring to previously published papers. The lipids are categorized into fatty acids, triglycerides, phospholipids, and terpenoid alcohols, along with their common names and structural annotations.

| Category | Subcategory | Compound name (Alternative name) | Short notation |
|---|---|---|---|
| Fatty Acids | Saturated Fatty Acids | Decanoic acid (Capric acid) | 10:0 |
| | | Undecanoic acid | 11:0 |
| | | Nonanoic acid | 9:0 |
| | | Octanoic acid | 8:0 |
| | Unsaturated Fatty Acids | cis-9-octadecenoic acid, (Oleic acid) | 18:1 |
| | | trans-9-octadecenoic acid (Elaidic acid) | 18:1 |
| | | cis-15-tetracosenoic acid (Nervonic acid) | 24:1 |
| | | cis-9-hexadecenoic acid (Palmitoleic acid) | 16:1 |
| | | cis-5-Dodecenoic acid | 12:1 |
| Triglycerides | Triacylglycerols (TG) | Tri-decanoyl-glycerol (Tricaprin) | TG (10:0/10:0/10:0) |
| | | Tri-hexadecanoyl-glycerol (Tripalmitin) | TG (16:0/16:0/16:0) |
